# Hepatic retromer is essential for systemic cholesterol homeostasis by regulating lysosomal cholesterol metabolism

**DOI:** 10.1101/2023.08.09.552611

**Authors:** Dyonne Y. Vos, Markus G. Barbosa, Andries H. Heida, A. Caillaud, Joël J. Tissink, Mirjam H. Koster, Marieke Smit, Nicolette Huijkman, Niels J. Kloosterhuis, A. Girardeau, Z. Begué-Racapé, Vincent W. Bloks, Rick Havinga, Marceline M. Fuh, Fulvio Reggiori, Muriel Mari, Ludger Scheja, Joerg Heeren, Folkert Kuipers, Jan Freark de Boer, Antoine Rimbert, Justina C. Wolters, Jan Albert Kuivenhoven, Cristy Verzijl, Bart van de Sluis

## Abstract

**Background:** Disturbed hepatic cholesterol homeostasis is associated with multiple diseases, including atherosclerotic cardiovascular disease and metabolic dysfunction– associated steatotic liver disease. The endo-lysosomal system is essential for cholesterol uptake and intracellular distribution, yet the mechanisms governing these processes remain incompletely understood. Here, we investigated the impact of hepatic VPS35, a subunit of the endosomal sorting complex retromer, on hepatocellular and whole-body cholesterol homeostasis.

**Methods:** We generated a liver-specific *Vps35* knockout mouse model (*Vps35*^HepKO^) and applied biochemical analyses, proteomics, and stable-isotope-labeled tracers to quantify critical processes of cholesterol metabolism. Human iPSC-derived liver organoids and CRISPR technology were used to translate our findings to humans. Mechanistic studies were performed in precision-cut liver slices from WT and *Vps35*^HepKO^ mice.

**Results:** Hepatic VPS35 deficiency led to an increase in endo-lysosomal degradative compartments and a marked reduction in specific lysosomal proteins, including lysosomal acid lipase (LAL), Scavenger Receptor Class B Member 2 (SCARB2), and Niemann-Pick type C1 (NPC1). Using pathway-specific inhibitors, we showed that VPS35 loss impairs the translation of SCABR2 and NPC1. Consistently, human iPSC-derived liver organoids lacking VPS35 also exhibited reduced expression of NPC1 and SCARB2 proteins. The decrease in these lysosomal proteins correlated with increased cholesterol levels in the plasma and liver of *Vps35*^HepKO^ mice. This was likely explained by the disrupted cholesterol trafficking through the endo-lysosomal system, delayed plasma cholesterol turnover, and upregulated cholesterol biosynthesis.

**Conclusion:** These findings uncover a previously unknown role for the hepatic retromer complex in maintaining systemic cholesterol homeostasis. Beyond its established function in endosomal cargo transport, we now demonstrate that retromer is also essential for lysosomal cholesterol handling. This role is mediated by regulating key lysosomal proteins involved in cholesterol metabolism, including LAL, NPC1, and SCARB2.

## Introduction

The liver is crucial for maintaining systemic cholesterol homeostasis, and its dysregulation is linked to various diseases, including atherosclerotic cardiovascular disease (ASCVD) and metabolic dysfunction–associated steatotic liver disease (MASLD)^1^. The endo-lysosomal system plays a central role in cholesterol metabolism, as it facilitates receptor-mediated uptake of low-density lipoprotein (LDL) and remnants, as well as the intracellular transport of cholesterol towards different compartments, such as plasma membrane (PM), endoplasmic reticulum (ER), and trans-Golgi network (TGN)^2,3^.

In lysosomes, the lipoprotein-derived cholesteryl esters (CE) are hydrolyzed by lysosomal acid lipase (LAL) to release fatty acids (FA) and free cholesterol (FC)^4^. Lysosomal FC egress is controlled by Niemann-Pick type C proteins (NPCs). The small luminal NPC2 protein binds FC and delivers it to the lysosomal membrane transporter NPC1^5^. Together, they enable the distribution of FC to the PM, ER, and TGN^3^. The scavenger receptor class B member 2 (SCARB2), also known as lysosomal integral membrane protein type 2 (LIMP2), acts in parallel with NPCs to mediate lysosomal cholesterol export^6^. In the ER, where FC levels are very low under normal conditions^7^, cholesterol is sensed by the sterol regulatory-element binding protein (SREBP)/SREBP cleavage-activating protein (SCAP) machinery to regulate cholesterol biosynthesis and LDL-cholesterol uptake in response to the cellular needs^8^.

The importance of the endo-lysosomal system in maintaining cholesterol homeostasis is well demonstrated by our recent studies and those of colleagues. For example, we have shown that the endosomal protein sorting complexes, such as the Wiskott-Aldrich syndrome protein and SCAR homolog (WASH) and the COMMD/CCDC22/CCDC93 (CCC) complexes, play a key role in regulating plasma cholesterol levels in mice, dogs, and humans^9–11^. These two complexes act together to facilitate the endosomal recycling of the LDL receptor (LDLR) and its family member LDLR-related protein 1 (LRP1), which are key for hepatic lipoprotein uptake (Figure S1A)^9–11^. Furthermore, in patients with lysosomal acid lipase deficiency (LAL-D, MIM #613497) or NPC diseases (MIM #257220, MIM #607625), cellular and systemic cholesterol homeostasis is also disturbed. These rare disorders are caused by loss-of-function mutations in *LIPA*, the gene encoding LAL, and *NPCs* genes, respectively^12,13^. LAL-D is characterized by the accumulation of CE in late endosomes and lysosomes, which mainly affects macrophages, the spleen, and the liver (MIM #613497). Complete loss of LAL in humans results in early-onset Wolman’s disease, which is fatal within the first year of life (MIM #620151)^14^. Milder mutations in LAL cause cholesteryl ester storage disease (CESD; MIM #278000), which has a later onset due to 5-10% residual LAL activity, resulting in a range of clinical phenotypes, including hypercholesterolemia, premature atherosclerosis, hepatosplenomegaly, and progressive liver disease^13,14^. NPC disease (MIM #257220, MIM #607625), characterized by FC accumulation in lysosomes of different organs, is primarily a neurodegenerative disorder^12^. However, it also leads to progressive liver disease associated with hepatosplenomegaly, cholestasis, fibrosis, and an increased risk of hepatocellular carcinoma^12,15^. To date, no cures are available for LAL-D nor NPC disease, and current treatments only alleviate symptoms^12,13^. Despite the well-described roles of LAL and NPCs in controlling cellular cholesterol homeostasis, the mechanisms regulating their levels and functions are ill-defined.

Recent studies have implied a role for the endosomal protein sorting machinery complex, retromer, in cholesterol homeostasis. Retromer consists of the vacuolar protein sorting VPS26, VPS29, and VPS35, and mainly localizes to endosomal membranes (Figure S1A, B). In concert with sorting nexins (SNXs), like SNX27, retromer regulates the endosomal sorting of an array of integral membrane cargo proteins and recycles these to the plasma membrane or the TGN for reuse^16–19^. Marquer *et al.* were the first to link retromer dysfunction to aberrant cholesterol homeostasis *in vitro*^20^. They showed that the transport of NPC2 from the TGN to the lysosome is mediated via the cation-independent (CI) mannose-6-phosphate receptor (M6PR). CI-M6PR is a retromer cargo^21,22^, and loss of retromer causes lysosomal cholesterol accumulation in HeLa cells due to impaired CI-M6PR retrograde trafficking^20^. Others showed that cholesterol accumulation in *in vitro* NPC1-deficiency models is associated with reduced retromer function^23^. Furthermore, *in vitro* studies have shown that retromer is essential for the recruitment of the WASH and CCC complexes to endosomes. However, a retromer-independent model for recruiting these two multiprotein complexes to the endosomes has also been postulated (Figure S1A, B)^24,25^.

Together, these data suggest a relation between retromer and cholesterol homeostasis, but the molecular connection and physiological relevance remain to be fully elucidated. In this study, we revealed a previously unknown role for hepatic retromer in lysosomal cholesterol metabolism, thereby providing novel insights into how the endo-lysosomal system contributes to maintaining whole-body homeostasis.

## Materials and Methods

### Animals

Mice carrying a *Vps35* allele flanked with *LoxP* sites (*Vps35*^fl/fl^; 129/Ola background)^26^ were crossed with a transgenic mouse line (C57Bl/6 background) expressing Cre recombinase under the control of the *Albumin* (*Alb*) gene promoter (#003574, The Jackson Laboratory), to generate a liver-specific *Vps35* knockout (*Vps35*^HepKO^) mouse model. Somatic ablation of *Vps35* in mouse livers was accomplished by a somatic CRISPR-mediated gene editing approach as described previously^27^. The sequence of *Vps35* sgRNA’s used is indicated in Figure S4C. To ablate *Vps35* in a *Ldlr* knockout background, mice expressing Cas9 specifically in the liver (#026556, The Jackson Laboratory)^28^ were crossed with *Ldlr* knockout mice (#002207, The Jackson Laboratory).

All animal experiments were approved by The Central Authority for Scientific Procedures on Animals (CCD), as well as by the Animal Welfare Body (IvD) of the University of Groningen (Groningen, the Netherlands). Male and female mice were used for the experiments as indicated in the results section. Mice were individually housed under a 12-h light-dark cycle and were fed *ad libitum* with standard laboratory diet (V1554-703, Ssniff Spezialdiäten GmbH), or cholesterol-enriched high-fat (HFC) diet (60% fat, 0.25% cholesterol; D14010701, Research Diets), and water. Body composition (lean, fat, and fluid mass) was measured after 12 weeks of HFC diet-feeding in unanesthetized mice, using nuclear magnetic resonance (NMR) MiniSpec (MiniSpec LF90 BCA-analyzer, Bruker).

Mice were sacrificed after 4 h of fasting. Blood was collected through cardiac puncture under anesthesia in EDTA-coated tubes. Plasma was collected after centrifugation at 1000 x *g* for 10 min at 4°C. Liver tissues were harvested and either snap-frozen in liquid nitrogen and stored at -80°C until further analysis, or fixed in 4% paraformaldehyde (PFA), embedded in paraffin, and cut into 4 μm sections for histological analysis.

### VLDL-TG production assay

After 4 h of fasting, mice were injected intraperitoneally with Poloxamer-407 (Sigma) solution in PBS (1 g per kg body weight) to block lipoprotein lipase (LPL) activity. Blood samples were taken from the tail vein at time points 0, 30, 60, 120, and 240 min after Poloxamer-407 injection. Blood samples were centrifuged at 1000 x *g* for 10 min at 4°C to collect plasma, which was used for triglyceride (TG) measurements. In addition, nascent VLDL particles (d<1.006 g/ml) were isolated from plasma collected at t=240 min by density gradient ultracentrifugation. Gradients were centrifuged overnight at 40000 rpm in a Beckman Coulter SW41 Ti rotor at 4°C. Concentrations of TGs, cholesterol, and phospholipids were measured, and ApoB levels were determined with targeted proteomics.

### Cholesterol flux measurements

Cholesterol fluxes were measured using a dual stable isotope tracer method as previously described.^29^ To determine intestinal cholesterol absorption, mice were administered cholesterol-D_5_ and cholesterol-D_7_, as follows: 0.3 mg cholesterol-D_5_ (Medical Isotopes Inc., Pelham, NH) was dissolved in 150 μL 20% Intralipid (Fresenius Kabi, Den Bosch, the Netherlands) and was injected intravenously. 0.6 mg cholesterol-D_7_ (Cambridge Isotope Laboratories, Inc., Andover, MA) was dissolved in 200 μL medium-chain TG oil and was administered by oral gavage. Bloodspots were collected from the tail vein prior to and at 3, 6, 9, 12, 24, 36, 48, 60, 72, 96, 120, 144, and 168 hours after administration for measurement of tracer enrichments. Standard pharmacokinetics were applied for fitting the enrichment curves using SAAM-II software (version 2.3, The Epsilon Group, Charlottesville, VA). The formula used for curve fitting was *f (t)* = *f_1_ e^-k1t^* + *f_2_ e^-k2t^* – *f_3_ e^-k3t^*.

To assess whole-body cholesterol synthesis using mass isotopomer distribution analysis (MIDA)^30,31^, mice received drinking water containing 2% [1-^13^C] acetate and bloodspots were collected at 0, 24, 33, 48, 57, 72, 81, 96, 105, and 120 hours. After the final time point, mice were sacrificed, and livers were collected for biochemical characterization.

### Histological analysis

Paraffin-embedded liver samples were cut into sections of 4 μm, which were stained with hematoxylin-eosin (H&E). Cell death was assessed by Terminal deoxynucleotidyl transferase-mediated dUTP nick-end labeling (TUNEL; S7100, Merck Millipore). Images were obtained with a Leica DM3000 microscope with a mounted Leica DFC420 camera. To quantify cell death, the number of TUNEL-positive cells was counted in ten high-power fields with 20x magnification for each liver sample of each experimental group. For each sample, the average of positive cells per field was calculated, which was used to calculate the average per field per experimental group.

### Scanning transmission electron microscopy (TEM)

For TEM analysis, small pieces of fresh liver (1-2 mm^3^) were fixed in 2% PFA and 2% glutaraldehyde in 0.1 M sodium cacodylate buffer (pH 7.4) at 4°C. Tissue processing, including staining with neodymium, sectioning, and image acquisition, was performed as previously described^32,33^. The images of the preparations are available upon request.

### Hepatic lipid extraction

Liver homogenates (15% w/v in PBS) were used for lipid extraction according to Bligh and Dyer^34^. 200 μl of liver homogenate was added to 600 μl of demi-water, mixed with 3 ml chloroform/methanol (1:2 v/v) in a glass tube, and incubated for 30 min. Then, 1.2 ml of water and 1 ml of chloroform were added, mixed, and samples were centrifuged at 500 x *g* for 10 min at room temperature (RT). The organic layer was transferred to a new glass tube, and the solvent was evaporated with nitrogen at 50°C. The dried lipids were dissolved in 1 ml chloroform, from which 400 µl was again dried and dissolved in 500 µl of 2% Triton X-100 in chloroform, and evaporated with nitrogen at 50°C. Then, 500 µl of water was added, and samples were incubated for 15 min at 37°C before further analysis for cholesterol and TG concentrations.

### Lipid measurements in plasma, isolated VLDL, and liver

Colorimetric assays were used to determine total cholesterol (TC), FC, TG, and phospholipid concentrations. Cholesterol enzymatic reagents were used to determine TC (#11489232, Roche; Cholesterol FS, DiaSys) and FC concentrations (#113609910930, DiaSys), with the cholesterol standard FS (DiaSys) as a reference. TG levels were determined with the Trig/GB kit (#1187771, Roche) or Triglycerides FS (DiaSys), using the Precimat Glycerol standard (Roche) as a reference. Concentrations of phospholipids were determined with the Phospholipid FS kit (DiaSys) and the Phospholipids Standard FS (DiaSys) as a reference.

Lipidomic analyses of liver samples were performed using the Lipidyzer™ Platform (SCIEX, Framingham, MA, USA), as previously described^35^.

### Fast-performance liquid chromatography (FPLC)

TC and TG content of the major lipoprotein classes (VLDL, LDL, and HDL) was determined using FPLC analysis as previously described^27^. Briefly, measurements were performed with a system containing a PU-4180 RHPLC pump and a UV-4075 UV-Vis detector (Jasco). Measurements were done in individual plasma samples when enough sample was available, or in pooled plasma samples as indicated. The samples, diluted in PBS, were loaded onto a Superose^TM^ 6 Increase 10/300 GL column (GE Healthcare) for separation of lipoproteins at a flow rate of 0.31 ml/min. An enzymatic reagent for cholesterol (Roche) was added through a second flow at a flow rate of 0.1 ml/min.

### Liver enzyme measurements in plasma

Concentrations of alanine aminotransferase (ALT) and aspartate aminotransferase (AST) in plasma were measured with a clinical chemistry analyzer (Cobas 6000, Roche Diagnostics), using standard reagents (Roche Diagnostics).

### Sucrose gradients

A 20% w/v homogenate was prepared with a piece of snap-frozen liver tissue in homogenization buffer (50 mM Tris-HCl [pH 7.4], 250 mM sucrose, 25 mM KCl, 5 mM MgCl_2_, 3 mM imidazole, supplemented with protease inhibitors [Roche]). To remove nuclei and debris, homogenates were centrifuged at 1000 x *g* for 10 min at 4°C. Protein concentrations were determined using the Bradford method (Bio-Rad). 1.5 mg of liver homogenates were loaded onto a 3.7 ml continuous 10% to 40% sucrose gradient, which was centrifuged for 16 h at 40.000 rpm in a Beckman Coulter SW55 Ti rotor. 370 μl fractions were collected from the top and were used for cholesterol measurements using the Amplex™ Red Cholesterol Assay Kit according to the manufacturer’s instructions (A12216, Thermofisher). For further analysis with untargeted proteomics, 30 μl of each fraction was mixed with 10 μl of 4x LDS sample buffer.

### Cell culture

VPS35-deficient cells were generated in a Cas9-expressing Hepa1-6 murine hepatoma cell line (kindly provided by Noam Zelcer^36^), as previously described^27^. Adenovirus to express sgRNAs to target *Vps35* was generated as previously reported ^9^; sgRNA sequences are listed in Supplemental Table 1. The cells were cultured in Dulbecco’s modified Eagle medium (DMEM) GlutaMAX (Gibco), supplemented with 10% fetal calf serum, 1% penicillin-streptomycin, and 1.25 μg/ml puromycin, at 37°C and 5% CO_2_. Subsequently, cells were used to assess LAL activity (further described below).

### Precision-cut liver slices

Precision-cut liver slices (PCLS) were prepared from mouse liver biopsy cores obtained using a 6 mm punch tool (KAI Medical), following the general methodology outlined by Groothuis et al.^37^. Liver tissue sections, approximately 250–300 µm thick and weighing ∼5 mg, were generated with a Krumdieck Tissue Slicer (Alabama Research and Development). Each slice was individually placed in a well of a 12-wells plate containing Williams E medium (US Biological Life Sciences, C17082259), supplemented with 25 nM glucose and 50 µg/ml gentamycin (Invitrogen). Cultures were maintained at 37°C under continuous agitation (90 rpm) and exposed to an 80% OL / 5% COL atmosphere for up to 24 h. Liver slices were treated with 25 nM bafilomycin A1 or 2.5 mM ammonium chloride for 16 h to inhibit lysosomal degradation. To suppress the formation and release of extracellular vesicles and exosomes, PCLS were treated with 5 µM GW4869 or 250 nM manumycin A1 for 16 h. To block proteasomal protein degradation, PCLS were treated with 5 µM or 2.5 µM MG132 for 16 or 22 h, respectively. Brefeldin A at a concentration of 10 µg/ml for 22 h, refreshed every 6 h, was used to inhibit the transport of proteins from the ER to the Golgi apparatus. After each respective incubation period, three liver tissue slices for each experimental condition were pooled in a single 1.5 ml microcentrifuge tube. Subsequently, protein extraction was performed as described below.

### Genome editing of human induced pluripotent stem cells (hiPSCs) and genotyping

The *VPS35*^-/-^ hiPS cell line was derived from the genomic edition of a control hiPS cell line reprogrammed from urinary progenitor cells and previously characterized^38,39^. Genomic editing was performed using the Alt-R™ CRISPR-Cas9 system (Integrated DNA Technologies) following a described procedure^40^. Two CRISPR guide RNAs (TACGAACTTGTACAGTATGC and GGTGTGCAACATCCCTTGAG) targeting exons 4 and 5 of *VPS35,* respectively, were used to delete 655 bp. After genome editing and clonal selection, hiPSC clones were genotyped by PCR and Sanger sequencing. Primers are listed in Supplemental Table 2.

### hiPSCs culture and differentiation into liver organoids

hiPSCs were cultured under hypoxia (4% O_2_, 5% CO_2_) on plates coated with 0.05 mg/ml Matrigel® (Corning) in StemMACS^TM^ iPS-Brew medium XF (Miltenyi) and passaged using the Gentle Dissociation reagent (Stem Cell Technologies). In-suspension liver organoids were generated following a modified protocol previously described^41^. hiPSCs were first dissociated and seeded in 24-wells AggreWell™400 plates (Stem Cell Technologies) at 1000 cells/well as recommended by the manufacturer, in StemMACS^TM^ iPS-Brew XF medium supplemented with 10 μM Y27632 (Cell Guidance Systems) to form aggregates (day -2). The next day, the media was carefully changed using StemMACS^TM^ iPS-Brew medium XF without disturbing the aggregates. On day 0, aggregates were collected using a 37μm reversible strainer (Stem Cell Technologies) and washed with complete RPMI 1640 media supplemented with 1X insulin containing B27, 1X NEAA and 1X GlutaMAX^TM^-I (Life Technologies). The aggregates were further resuspended in a vented Erlenmeyer flask in complete RPMI medium supplemented with 4 μM CHIR99021 (BOC Sciences) to induce mesendoderm formation and incubated at 37°C under agitation. All subsequent steps were performed under orbital shaking at 70 rpm. On day 1 after initiation of the differentiation, the media was changed to complete RPMI medium without any other molecule to pattern aggregates towards definitive endoderm. From day 2 to 6, aggregates were directed towards hepatic endoderm with medium changes every 2 days using knockout DMEM supplemented with 20% Knockout serum replacement (KSR), 1X NEAA, 1X GlutaMAX^TM^-I (Life Technologies) and 1% DMSO and 100 μM 2-mercaptoethanol (Sigma). From day 7 to 19, liver organoids were cultured in liver maturation medium with medium changes every 2 to 3 d using hepatocyte culture medium (HCM) without EGF (Lonza) supplemented with 100 nM dexamethasone (Sigma) and 100 nM N-hexanoic-Tyr-Ile-(6) aminohexanoic amide (Active Peptide). Organoids were harvested on day 20 for further analysis. Cellular pluripotency was assessed by measuring the expression of *OCT4*, *NANOG*, *SOX2 and ALB, HNF4, AFP* as liver-like specific markers (Supplemental Table 2). RNA was extracted from hiPSCs and liver organoids using the NucleoSpin Tissue Purification Kit (Macherey-Nagel). Reverse transcription of RNA to cDNA was performed using the high-capacity cDNA reverse-transcription kit (Applied Biosystems). qPCR studies were performed in triplicate for hiPSCs and quadruplicate for liver organoids using the Power SYBR® Green PCR Master Mix on the QuantStudio^5^ system with automatic threshold calculation (Applied biosystems^TM^).

### Isolation of liver plasma membrane

Subcellular fractionation was obtained by using the Mem-PER Plus Membrane Protein Extraction kit (Life Technologies). Liver tissue weighing 20–40 mg was placed in a 5 ml microcentrifuge tube. 4 ml of cell wash solution was added to the tube, followed by a brief vortex. The wash solution was discarded. The tissue was transferred to a 2 ml tissue grinder. 1 ml of permeabilization buffer was added, and the tissue was homogenized with 6–10 strokes until an even suspension was achieved. An additional 1 ml of permeabilization buffer was added to the homogenate, which was then transferred to a new tube and incubated at 4°C for 10 min with constant mixing. The permeabilized suspension was centrifuged at 16,000 × g for 15 min at 4°C to obtain a pellet. The supernatant containing the cytoplasmic fraction was removed and transferred to a new tube. The pellet was resuspended in 1 ml of solubilization buffer by pipetting up and down to create a homogeneous suspension. The sample was incubated for 30 min at 4°C with constant mixing. Following incubation, the tubes were centrifuged again at 16,000 × g for 15 min at 4°C. The supernatant containing solubilized membrane and membrane-associated proteins was collected in a new tube. The prepared samples were stored at -80 °C or used directly for downstream analysis.

### In vivo biotinylation assay

*In vivo* biotinylation was performed as reported previously with minor modifications^42^. Mice were anesthetized via intraperitoneal injection of ketamine (7 5mg/kg), xylazine (20 mg/kg), and acepromazine (3 mg/kg). Following anesthesia, transcardial perfusion was performed with 10 ml of pre-warmed PBS containing 10% dextrane 40 for 10 min at a flow rate of 1 ml/min, and subsequently, with 15 ml of pre-warmed biotinylation buffer containing non-permeable, cleavable 1 mg/ml EZ Link Sulfo-NHS-SS-Biotin (Thermofisher, #21331) in PBS containing 10% dextrane 40 for 10 min at a flow rate of 1.5 ml/min. To inactivate the EZ Link Sulfo-NHS-SS-Biotin, mice were perfused with pre-warmed 15 ml of inactivation buffer, containing 50 mM Tris/HCl and 10% dextrane 40 in PBS, at a flow rate of 1.5 ml/min. In parallel, the liver surface was sprinkled with inactivation sprinkling buffer containing 50 mM Tris/HCl in PBS and then harvested and snap-frozen. For the pulldown of biotinylated surface proteins, livers were homogenized using 15 strokes of a Dounce homogenizer in IP lysis buffer (25 mM Tris/HCl [pH 7.4], 150 mM Nacl, 1mM EDTA, 1% NP-40, 5% glycerol) supplemented with phosphatase and protease inhibitors, followed by sonification for 20 s. Equal amount of protein for each sample was incubated with pre-washed NeutrAvidin Agarose Resins (Thermofisher, #29200). Afterwards, NeutrAvidin Agarose Resins were washed three times and biotinylated proteins were eluted with 2x SDS sample buffer containing 0.25 M DTT, boiled for 5 min at 95°C and analyzed using immunoblotting as described below.

### Gene expression analysis

A piece of snap-frozen liver tissue was homogenized in QIAzol Lysis Reagent (Qiagen). RNA was isolated by chloroform extraction, precipitated with isopropanol, washed with ethanol, and dissolved in RNase/DNase-free water. 2 μg of RNA were used as input for cDNA synthesis, and 20 ng of cDNA was used for quantitative reverse transcription PCR with the FastStart SYBR Green Master (Roche) and the QuantStudio 7 Flex Real-Time PCR System (Applied Biosystems). The qRT-PCR program was as follows: 50°C/2 min; 95°C/10 min; 40 cycles with 95°C/15 s; and 60°C/1 min. Gene expression was calculated using the ΔΔCT method using QuantStudio Real-Time PCR Software (Applied Biosystems), with *Ppia* expression as an internal control. Primer sets used for gene expression analysis are listed in Supplemental Table 3.

For RNA sequencing analysis, 6 representative liver samples were selected, and total RNA was isolated with the RNAeasy plus kit (#74134, Qiagen) according to the manufacturer’s instructions. Quantity and quality were determined using nanodrop and gel electrophoresis, respectively. Library preparation, sequencing, and analysis were done by Novogene Co. Ltd. Europe. A heatmap was constructed using the z-score calculated from the FPKM values, to visualize the expression of genes involved in cholesterol biosynthesis.

### Immunoblotting

Liver and cell samples were homogenized in NP-40 buffer (0.1% Nonidet P-40, 0.4 M NaCl, 10 mM Tris-HCl [pH 8.0], and 1 mM EDTA) supplemented with 1 mM DTT and protease and phosphatase inhibitors (Roche). Protein concentrations were measured with the Bradford assay (Bio-Rad), and samples were diluted in 4x sample buffer. Samples were boiled for 5 min at 95°C prior to loading on gel, except for samples used to visualize NPC1 and SCARB2. Proteins were separated with SDS-PAGE and subsequently transferred to Amersham Hybond-P PVDF Transfer Membrane (RPN303F, GE Healthcare). Membranes were blocked in 5% milk in TBS buffer containing 0.01% Tween 20 and incubated with the antibodies listed in Supplemental Table 3. Proteins of interest were visualized and quantified with the ChemiDoc XRS+ System and Image Lab software version 5.2.1 (Bio-Rad).

### Targeted proteomic analysis

To determine protein levels of ApoB (ApoB48 and ApoB100) in plasma and isolated VLDL samples (isolated as described above), we used a targeted proteomic assay as previously described.^27^ For plasma samples, we selected 5 representative samples from each group. In-gel digestion was performed on 1 μl of plasma and 22.5 μl of VLDL samples, and the amount of isotopically-labeled standards added for the quantification was optimized for each sample type. LC-MS analyses and data processing and quantification were done as previously reported.^9^

### Discovery-based (untargeted) proteomic analysis

Discovery-based proteomics was used to determine relative protein concentrations in whole liver samples and sucrose-fractionated liver samples. For whole liver samples, 6 representative samples were selected from each group. In-gel digestion was performed on 50 μg of liver lysates or 30 μl from the collected sucrose fractions as described previously^9^.

For the whole liver lysates, the discovery-based proteomics analyses (Label Free Quantification) were performed as described previously^43^. A rerun of the whole liver lysates was done with a digestion based on the published SP3 digestion protocol^44^ using the discovery-based proteomics settings similar to the sucrose gradient samples, to detect SOAT2. For the SP3 digestion protocol, 10 μg of total protein was diluted in 100 mM ammonium bicarbonate to a final volume of 25 μl as starting material. Reduction was performed by adding 10 μl 10 mM dithiothreitol and incubation for 30 min at 57 °C. Alkylation was done with 10 μl of 30 mM iodoacetamide for 30 min at RT under aluminum foil. The proteins were bound to 10 μl prewashed beads (1:1 mix of hydrophilic and hydrophobic Cytiva SeraMag beads, Fisher Scientific), and the samples were diluted to 50% acetonitrile with 55 μl acetonitrile. After removal of the acetonitrile, the beads were washed twice with 200 μl 80% v/v ethanol, once with 180 μl acetonitrile, and resuspended in 40 μl 2.5 ng/μl trypsin (V5111 sequencing grade modified trypsin, Promega) for overnight digestion at 37 °C. The digestion was stopped by addition of 10 μl 1% v/v formic acid and an equivalent of 1 μg was loaded on the Evosep tips for the untargeted proteomics analyses.

For the sucrose gradient samples, the discovery mass spectrometric analyses were performed on a quadrupole orbitrap mass spectrometer equipped with a nano-electrospray ion source (Orbitrap Exploris 480, Thermo Scientific). Chromatographic separation of the peptides was performed by liquid chromatography (LC) on an Evosep system (Evosep One, Evosep) using a nano-LC column (EV1137 Performance column 15 cm x 150 µm, 1.5 µm, Evosep; buffer A: 0.1% v/v formic acid, dissolved in water, buffer B: 0.1% v/v formic acid, dissolved in acetonitrile). Ten percent of the sucrose fraction digests were injected and separated using the 30SPD workflow (Evosep). The mass spectrometer was operated in positive ion mode and data-independent acquisition mode (DIA) using isolation windows of 16 m/z with a precursor mass range of 400-1000, switching the FAIMS between CV-45V and -60V with three scheduled MS1 scans during each screening of the precursor mass range.

Raw LC-MS data were processed with Spectronaut (16.0.220606) (Biognosys) with the standard settings of the directDIA workflow except that quantification was performed on MS1 with a mouse SwissProt database (www.uniprot.org, 17021 entries). For data quantification of total livers, local normalization was applied, and the Q-value filtering was set to the classic setting without applying imputation. With these data, z-scores were calculated and used to construct a heatmap to visualize the relative expression of lysosomal proteins. Alternatively, data were used to express relative SOAT2 levels. For sucrose fractionated samples, the Q-value filtering was set to the classic setting without applying normalization or imputation, and the data were used to visualize the expression of proteins in each sucrose fraction, relative to the total protein expression in all fractions.

### LAL activity assay

LAL activity was determined in control and VPS35-deficient Hepa1-6 cells using the LysoLive™ Lysosomal Acid Lipase Assay Kit (ab253380, Abcam) according to the manufacturer’s instructions for adherent cells. Cells were incubated with LysoLive^TM^ LipaGreen^TM^ for 6 h in serum-free medium prior to reading cells with flow cytometry. Cells were analyzed with the BD FACSVerse™ (BD, Biosciences). Data was processed with Kaluza software (Beckman Coulter).

### Immunofluorescence and cholesterol staining

Primary hepatocytes were plated onto collagen-coated glass coverslips (Corning, 11563550). After 24 h, cells were briefly washed with PBS at RT, followed by fixation in 4% PFA at RT for 10 min. Cells were then washed twice with PBS at RT for 5 min each and maintained in PBS. Permeabilization was performed by immersing the coverslip in liquid nitrogen for 5 s, followed by immediate transfer back into PBS. Samples were then incubated with blocking solution (1% human serum in PBS) at RT for 30 min under gentle agitation. Cholesterol in cell membranes was visualized with GFP-D4 probe (1:100 in PBS containing 1% human serum) for 1 h at RT^45^. After a brief PBS wash, cells were fixed again with 4% PFA for 10 min at RT and washed with PBS at RT for 5 min. Non-specific binding was further blocked with 1% human serum in PBS for 30 min. Cells were then incubated with rabbit anti-LAMP2A antibody (1:200, Abcam, ab18528) overnight at 4°C in a humidified chamber. Following three PBS washes at RT, cells were incubated with Cy3-conjugated donkey anti-rabbit IgG secondary antibody (1:300, Jackson ImmunoResearch, 711-165-152) for 1 h at RT. Samples were washed three additional times in PBS. Coverslips were mounted using VECTASHIELD® Antifade Mounting Medium with DAPI (VectorLabs, H-1200-10). Imaging was performed using a confocal laser scanning microscope.

### Statistical analysis

Unless indicated otherwise, data are presented as mean ±SEM. Statistical analyses were performed with GraphPad version 9.1 (GraphPad software). Differences between two groups were determined with unpaired two-tailed Student’s *t* test. For comparing two variables between groups, two-way ANOVA with Tukey’s multiple comparison test was performed. In all cases, P values of < 0.05 were considered statistically significant.

## Results

### Liver-specific depletion of VPS35 destabilizes retromer without affecting other endosomal sorting complexes

To examine the physiological role of retromer in cholesterol homeostasis, we generated a liver-specific *Vps35* knockout (*Vps35*^HepKO^) mouse model. Using immunoblotting, we confirmed the ablation of *Vps35* in the livers of *Vps35*^HepKO^ mice (Figure 1A). In line with previous studies^46,47^, loss of *Vps35* strongly decreased the protein levels of the other retromer subunits VPS26 and VPS29 by 91% and 83%, respectively (Figure 1A,B). Retromer shares its subunit VPS29 with retriever, which consists of VPS26C, VPS29, and VPS35L (Figure S1B), and shows structural similarities with retriever^48^. Retriever is physically and functionally connected with the COMMD1/CCDC22/CCDC93 (CCC) complex (Figure S1B)^27,48^, and therefore, we also determined the protein levels of VPS35L and the CCC components CCDC22, CCDC93, and COMMD1. Immunoblotting shows that hepatic loss of VPS35 did not affect the levels of these proteins (Figure S2A). To get further insight into the complex formation of the retriever, CCC, and WASH complexes, we determined the relative distribution of the components of these multiprotein complexes within the endo-lysosomal system, using sucrose density gradient fractionation of homogenates from *Vps35*^HepKO^ and control mouse livers. We analyzed the proteome in all fractions by protein mass spectrometry and detected the retromer subunits mainly in fractions 5-7 in control livers (Figure S2B) and confirmed the marked reduction of the subunits upon *Vps35* ablation (Figure S2B). The components of the retriever, CCC, and WASH complexes were mainly present in fractions 9-14, and loss of VPS35 did not affect their relative distribution in the gradient (Figure S2B). These data indicate that hepatic *Vps35* ablation specifically impacts retromer without affecting the formation of retromer-associated protein complexes, such as retriever, CCC, and WASH complexes.

**Figure 1.**
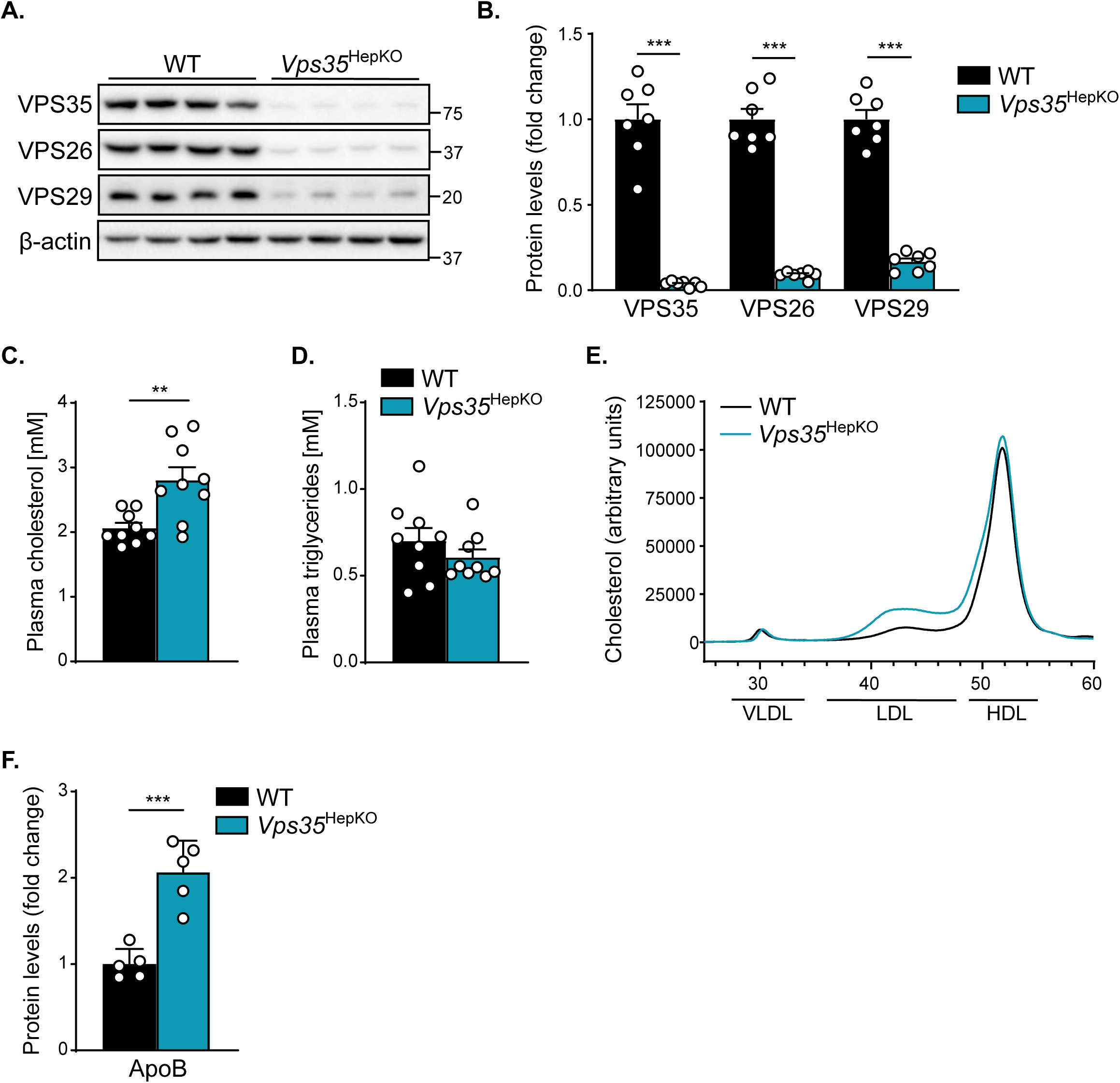
Liver-specific VPS35-deficient mice are hypercholesterolemic. (**A**) Hepatic protein levels of the retromer subunits VPS35, VPS26, and VPS29 were determined by immunoblotting (n=4). (**B**) Quantification of hepatic protein levels of the retromer subunits, as determined by immunoblotting (n=7). (**C**) Plasma cholesterol and (**D**) TG levels in WT and *Vps35*^HepKO^ male mice (n=9). (**E**) Cholesterol levels in plasma fractionated by FPLC (n=7). (**F**) ApoB levels in plasma, determined by targeted proteomics (n=5). Data are presented as mean ± SEM; ** p < 0.01 and *** p < 0.001.

### Hepatic VPS35 deficiency increases plasma cholesterol levels

*Vps35*^HepKO^ male mice were born at the expected Mendelian ratio, but the body and liver weights of these animals were slightly higher compared to their wild-type (WT) littermates (Figure S3A, B). To determine the effect of hepatic *Vps35* depletion on plasma lipid levels, we measured cholesterol and TG concentrations in plasma samples. We found that *Vps35*^HepKO^ mice showed a 36% increase in plasma cholesterol levels compared to control mice (Figure 1C). Plasma TG levels were, however, not different between control and *Vps35*^HepKO^ mice (Figure 1D). Fast protein liquid chromatography (FPLC) separation showed that the elevated plasma cholesterol levels in *Vps35*^HepKO^ mice were mainly caused by an increase in LDL cholesterol (Figure 1E), which was supported by a two-fold increase in plasma levels of ApoB (Figure 1F).

### Hepatic ablation of VPS35 specifically impairs LRP1 function

Retromer is physically associated with the WASH complex, a complex that acts together with the CCC complex to recycle LDLR and LRP1 from endosomes back to the PM (Figure S1A, B)^9–11^. Since hepatic depletion of either WASH or CCC reduces surface LDLR and LRP1 levels, accompanied by elevated plasma LDL cholesterol levels^9–11^, we studied the hepatic levels of both receptors in *Vps35*^HepKO^ and control mice. We found that loss of VPS35 resulted in a 50% increase in hepatic LDLR protein levels and a 40% decrease in LRP1 levels (Figure 2A,B). These changes in protein levels could not be explained by their respective mRNA levels (Figure S4A). To further explore the effect of VPS35 deficiency on the localization of LDLR and LRP1 at the PM, we performed an *in vivo* PM biotinylation experiment. VPS35 deficiency reduced the protein levels of LRP1 at the PM but not LDLR levels (Figure 2C), indicating that retromer facilitates the endosomal recycling of LRP1 but not LDLR. The decrease in surface LRP1 levels, without affecting LDLR PM levels, was confirmed by isolating PM fractions from WT and *Vps35*^HepKO^ livers (Figure 2D). This specific adverse effect on LRP1 function is supported by the profound increase in hepatic LDLR levels (Figure 2A), as previous work in mice showed that impaired functioning of LRP1 is compensated by an increase in LDLR expression^49^. However, this increase in LDLR levels could also be explained by a decrease in the plasma levels of proprotein convertase subtilisin-kexin type9 (PCSK9). PCSK9 is secreted by the liver and targets LDLR for proteolysis^50,51^, and a recent study showed that loss of VPS35 decreases Sortilin-mediated PCSK9 expression^52^. To this end, we assessed the plasma PCSK9 by proteomics analysis and found that PCSK9 levels were significantly increased in *Vps35*^HepKO^ mice compared to control mice (Figure S4B).

**Figure 2.**
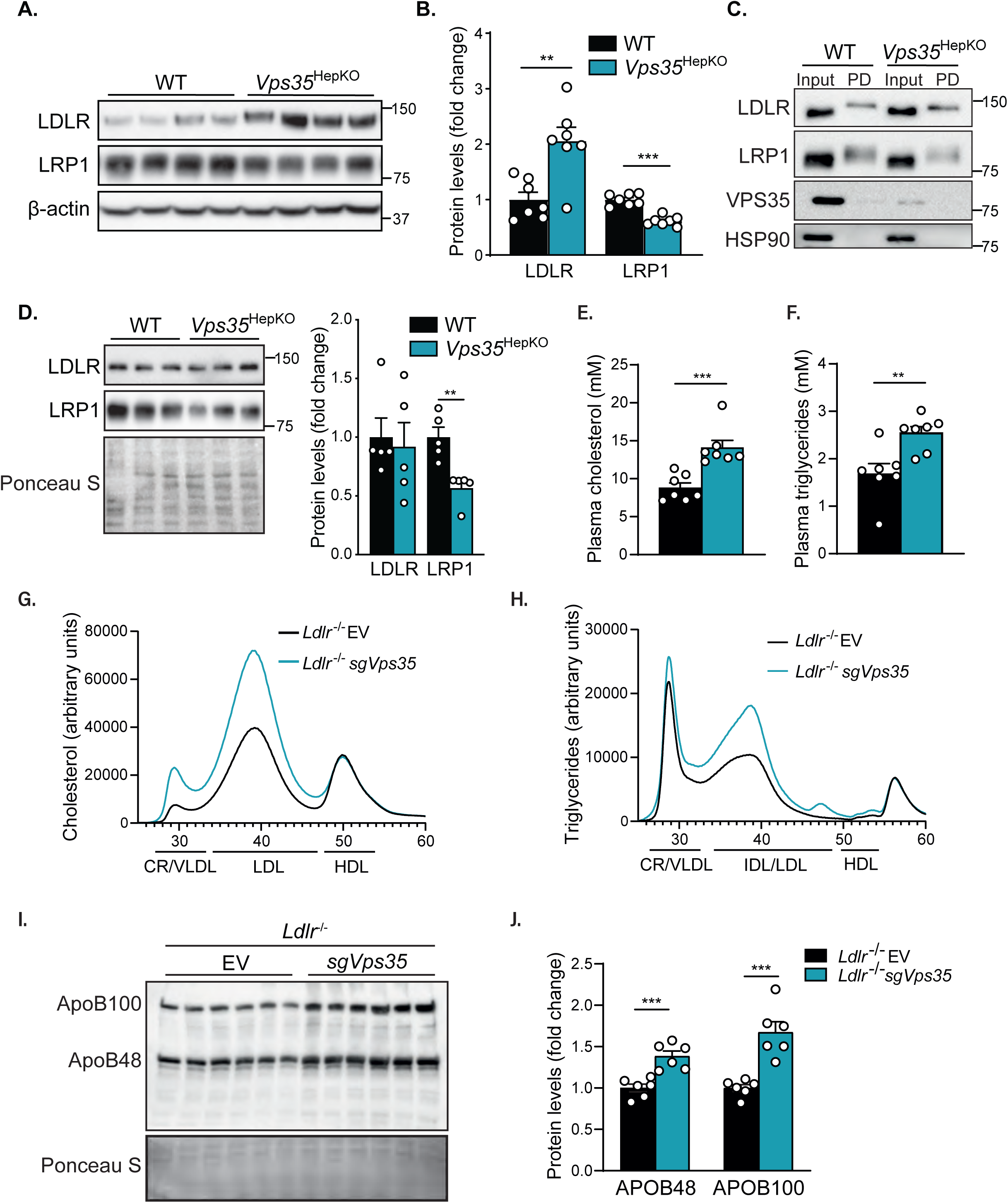
Hepatic VPS35 is required for LRP1 functioning to clear apoB-containing lipoprotein particles. (**A**) Immunoblot of LDLR and LRP1 in livers of WT and *Vps35*^HepKO^ male mice (n=4). (**B**) Quantification of LDLR and LRP1 protein levels (n=7), as determined by immunoblotting. (**C**) *In vivo* cell surface biotinylation was performed in WT and *Vps35*^HepKO^ male mice. LDLR and LRP1 were detected in total liver lysates (input, 0.5%) and the isolated cell surface protein fraction by immunoblotting. (**D**) LDLR and LRP1 at the plasma membrane after enrichment of the plasma membrane. Quantification of LDLR and LRP1 levels at the plasma membrane, as determined by immunoblotting (n=5). (**E**) Plasma cholesterol and (**F**) TG levels in WT and sg*Vps35*; LDLR-deficient male mice (n=9). (**G**) Cholesterol and (**H**) TG levels in plasma fractionated by FPLC of WT and sg*Vps35*; LDLR-deficient male mice (n=9). (n=7). (**I**) Immunoblot of apoB in plasma of WT and sg*Vps35*; LDLR-deficient male mice (n=6). (**J**) Quantification of APOB48 and APOB100 protein levels (n=6) determined by immunoblotting. Data are presented as mean ± SEM; ** p < 0.01 and *** p < 0.001.

To directly assess the role of VPS35 in LRP1 function, we ablated hepatic *Vps35* in *Ldlr* knockout (*Ldlr*^-/-^) mice by somatic CRISPR/Cas9-mediated gene editing (Figure S4C, D). Four weeks after injection of adeno-associated virus (AAV) encoding sgRNAs targeting *Vps35* in *Ldlr*^-/-^ mice (Figure S4E, F), we found that hepatic loss of retromer significantly increases plasma cholesterol and TG levels compared to control *Ldlr*^-/-^ mice (Figure 2E, F). These increases were caused by elevated chylomicron remnant/VLDL and LDL levels, as determined by FPLC separation (Figure 2G, H), which was supported by a significant increase in plasma levels of ApoB48 and ApoB100 (Figure 2I, J). This phenotype shows strong similarities with the lipid phenotype seen in mice deficient for LDLR and hepatic LRP1^49^. Taken together, these data indicate that retromer is required for the endosomal transport of LRP1, but not that of LDLR.

### Vps35^HepKO^ mice accumulate both free and esterified cholesterol in their livers

To better understand how hepatic loss of retromer affects plasma cholesterol levels, we studied the effect of hepatic *Vps35* depletion on hepatic lipid levels. Hepatic TC content in *Vps35*^HepKO^ mice was significantly increased compared to control mice (Figure 3A), while TG concentrations tended to decrease (Figure 3B). The increase in TC was caused by higher FC as well as cholesterol esters (CE) (Figure 3A). Using lipidomics, we confirmed the increase in CE (+46%) and found a significant 26% decrease in hepatic TG levels (Figure 3C). In addition, we observed changes in other lipid species, including the sphingolipids sphingomyelin (SM) (-14%), dihydroceramide (DCer) (+31%), and lactosylceramide (LCer) (+193%), and the phospholipid lysophophatidylethanolamine (LPE), a minor component of cell membranes, which was elevated by 67% (Figure 3D).

**Figure 3.**
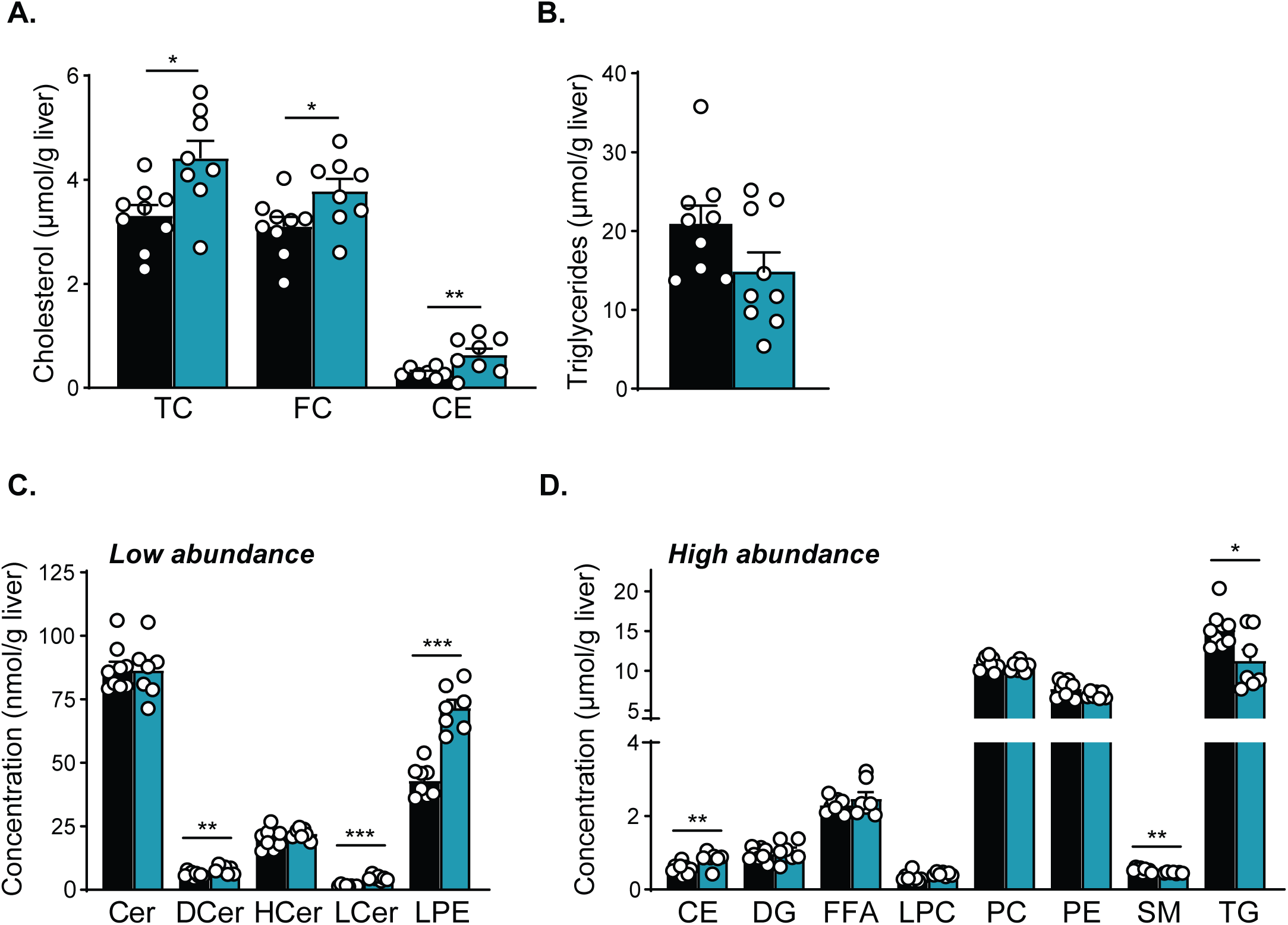
Hepatic *Vps35* depletion increases hepatic cholesterol content in mice. (**A**) Hepatic cholesterol (n=7-9) and (**B**) TG (n=9) levels in WT and *Vps35*^HepKO^ male mice. (**C, D**) Concentrations of lipid classes in livers of WT and *Vps35*^HepKO^ male mice, determined by lipidomics analysis (n=7-9). Data are presented as mean ± SEM; * p < 0.05, ** p < 0.01 and *** p < 0.001. *Abbreviations: ALT = alanine transaminase; AST = aspartate transaminase; CE = cholesterol ester; Cer = ceramide; DCer = dihydroceramide; DG = diacylglycerol; FC = free cholesterol; FFA = free fatty acid; HCer = hexosylceramide; LCer = lactosylceramide; LPC = lysophosphatidylcholine; LPE = lysophosphatidylethanolamine; PC = phosphatidylcholine; PE = phosphatidylethanolamine; SM = sphingomyelin; TC = total cholesterol; TG = triacylglycerol*.

The increase in hepatic cholesterol content was accompanied by elevated plasma levels of the liver enzymes ALT and AST (Figure S5A), suggesting that hepatic loss of VPS35 induces liver damage. However, histological examination using hematoxylin and eosin (H&E) and TUNEL staining did not reveal overt liver damage or hepatocellular apoptosis, respectively (Figure S5B, C). In addition, we found an increase in the expression of the monocyte/macrophage marker *Cd68* and monocyte chemoattractant protein 1 (*Mcp1*), but no significant changes in the expression of other inflammatory markers, such as *Il1*β, *Il6*, *Tnf*, and *F4/80* (Figure S5D). These findings show that hepatic retromer is essential to maintain hepatic and plasma cholesterol homeostasis, and its loss leads to mild liver damage.

### Hepatic VPS35 loss decreases the levels of a subset of lysosomal proteins involved in intracellular cholesterol transport

Hepatic lysosomes play an essential role in hepatic and plasma cholesterol homeostasis, and previous *in vitro* work has shown that VPS35 deficiency causes changes in late endosomal/multivesicular body (MVB) and lysosomal structure, and aberrant lysosomal functions ^46,47,53–55^. For this reason, we analyzed the ultrastructure of MVBs and the degradative compartments, i.e., lysosomes and autolysosomes/amphisomes, in VPS35-deficient hepatocytes using scanning electron microscopy. We found that the number of degradative compartments and MVBs was significantly increased in these cells compared to the WT controls (Figure 4A, B, C), suggesting that VPS35 depletion negatively affects the endo-lysosomal system. To obtain further insight into this, we examined the levels of late endosomal/lysosomal proteins by proteomics. We found that multiple lysosomal proteins were unaffected by the loss of VPS35 (Figure 4D), while some were significantly increased, including the V-ATPase subunits VATC1 and VATB2, cathepsin B, and NPC2, or decreased, including RAB7A, TPP1, RISC, cathepsin A/PPGB, and LAL (Figure 4D). Interestingly, a marked reduction was also observed in SCARB2 levels. The lysosomal protein NPC1 was not identified in the proteomic approach, but because of its essential role in lysosomal cholesterol export, we determined its levels by immunoblotting together with those of SCARB2. Surprisingly, and in contrast to LAMP1 used a control, NPC1 levels, as those of SCARB2, were also strongly decreased in *Vps35*^HepKO^ livers compared to controls (Figure 4E, F). In line with its decreased protein levels, we found that LAL activity was significantly reduced in *Vps35* KO cells compared to controls (Figure 4G). The lower levels of NPC1, SCARB2, and LAL were not due to changes in mRNA levels (Figure S6A), suggesting that they were the consequence of a post-transcriptional event. A reduction in NPC1 and SCARB2 was also observed in adult female *Vps35*^HepKO^ mice (Figure S6B), indicating that the effect of VPS35 deficiency is not sex specific. LAL was not examined due to the lack of a commercially available antibody against this enzyme. To assess whether the reduced levels of these lysosomal proteins result from compensatory adaptations to chronic retromer deficiency starting at birth, we deleted VPS35 in adult mice (8-9 weeks old) using a somatic CRISPR/Cas9-mediated gene editing approach (Figure S4B). Two weeks after *Vps35* ablation (Figure S6C), we sacrificed the mice and examined NPC1 and SCARB2 levels. Acute hepatic ablation of VPS35 in adult mice strongly reduced NPC1 and SCARB2 levels compared to the control mice (Figure S6D), suggesting that the effect of retromer deletion on these specific lysosomal proteins is likely not an adaptive mechanism. Collectively, these data show that hepatic loss of retromer, in males and females, has a strong negative effect on the expression of a subset of lysosomal proteins, including key proteins required for cholesterol metabolism and transport, such as LAL, NPC1, and SCARB2.

**Figure 4.**
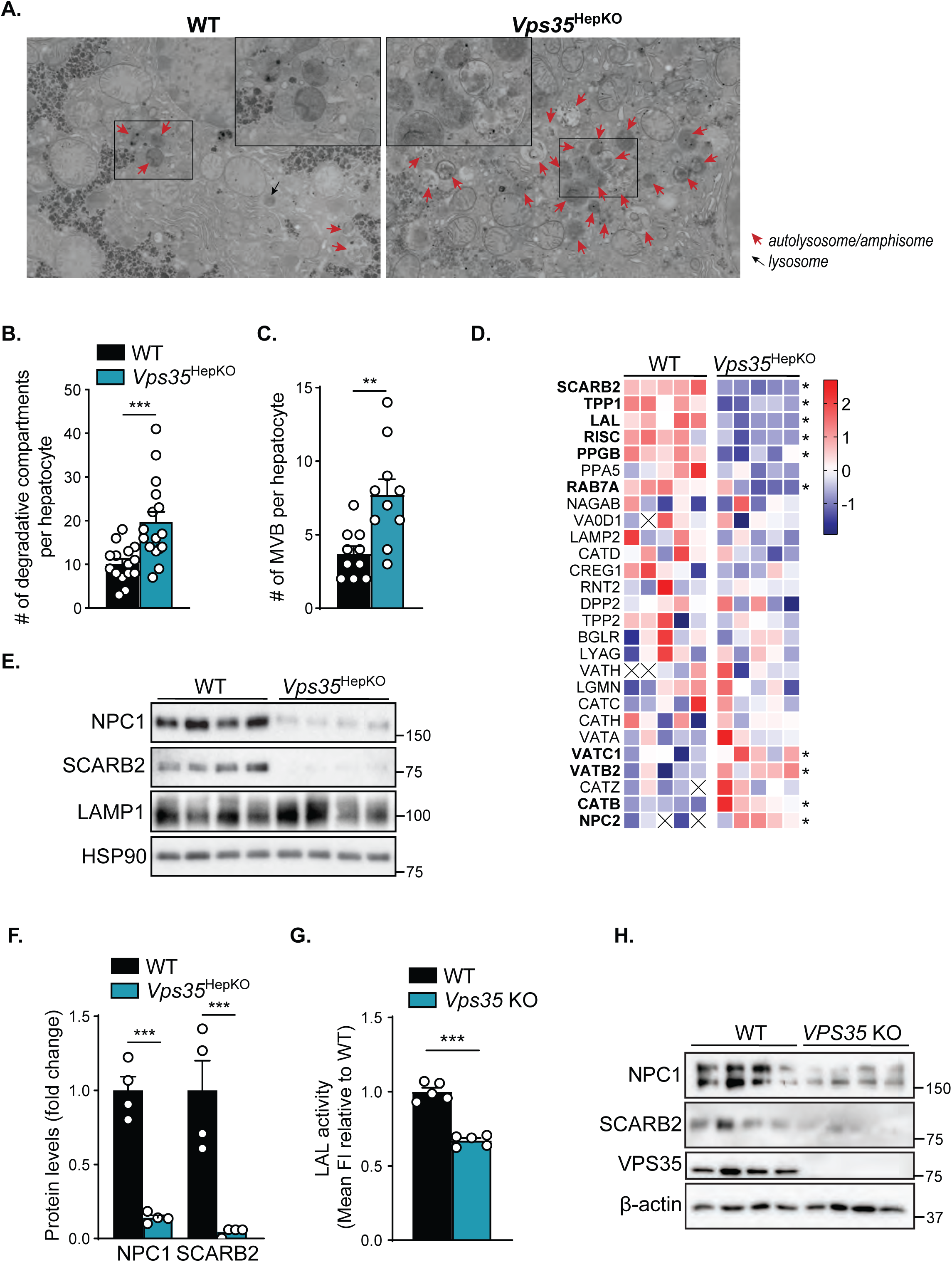
Hepatic loss of VPS35 affects the levels of various lysosomal proteins, including cholesterol transporters. (**A**) Scanning transmission electron microscopic images of livers of WT and *Vps35*^HepKO^ male mice, where amphisomes, autolysosomes, and lysosomes are indicated. Full datasets are available upon request. (**B**) Quantification of degradative compartments and (**C**) multivesicular bodies (MVB) in WT and *Vps35*^HepKO^ hepatocytes. (**D**) Heatmap showing z-scores of protein levels of LE/lysosomal proteins, presented as protein names, calculated from raw counts measured with untargeted proteomics (n=5); significantly changed proteins are indicated in bold and with *. (**E**) Immunoblot of NPC1, SCARB2, and LAMP1 in WT and *Vps35*^HepKO^ male livers (n=4). (**F**) Quantification of NPC1 and SCARB2 protein levels (n=4), as determined by immunoblotting (**E**). (**H**) LAL activity, expressed as mean fluorescence intensity, in WT and VPS35-KO Hepa1-6 cells, determined by FACS analysis (n=5 from two independent experiments). (**I**) NPC1 and SCARB2 protein levels in WT and VPS35-deficient Human iPSC-derived liver organoids (WT and VPS35 knockout iPSC clones were differentiated into liver organoids, data presents 4 independent differentiation experiments). Unless indicated otherwise, data are presented as mean ± SEM; ** p < 0.01 and *** p < 0.001.

### Loss of VPS35 in hiPSC-derived liver organoids decreases NPC1 and SCARB2

To translate our findings to humans, we ablated *VPS35* in human-induced pluripotent stem cells (hiPSCs) using CRISPR/Cas9-mediated gene editing approach. We successfully targeted VPS35, resulting in the complete absence of VPS35 protein expression in hiPSCs (Figure S7A). We subsequently differentiated control and VPS35-deficient hiPSCs into liver organoids. The success of the differentiation process was confirmed by analyzing mRNA levels of pluripotency and hepatocyte markers (Figure S7B, C). We then assessed NPC1 and SCARB2 expression by immunoblotting and found that the loss of VPS35 in hiPSC-derived liver organoids significantly decreased the expression of these lysosomal proteins (Figure 4I), without affecting their mRNA levels (Figure S7D). Again, we were unable to study the protein expression of LAL due to the lack of antibodies. In summary, and consistent with our findings in mouse livers, human VPS35 plays an essential role in maintaining the levels of NPC1 and SCARB2 in hepatocytes.

### Hepatic VPS35 deficiency increases cholesterol content of intracellular compartments

To get insight into the relative distribution of NPC1, SCARB2, LAL, and cholesterol within intracellular organelles, including endo-lysosomal compartments (Figure S2B), we fractionate mouse liver homogenates on sucrose density gradients and examined each fraction by proteomics. We found that NPC1, SCARB2, and LAMP1 were primarily present in fractions 13-15 (Figure 5A). LAMP2 was also present in fractions 13-15 but was mainly detected in fractions 3-5, where it co-sedimented with LAL. NPC1, SCARB2, and LAL levels were decreased in most of the fractions derived from VPS35-deficient livers. Although the LAMP1 levels were unchanged, its relative distribution was shifted from fractions 13-15 to fractions 10-12 upon *Vps35* ablation (Figure 5A), which might indicate that the morphology of LAMP1-positive degradative compartments is changed.

**Figure 5.**
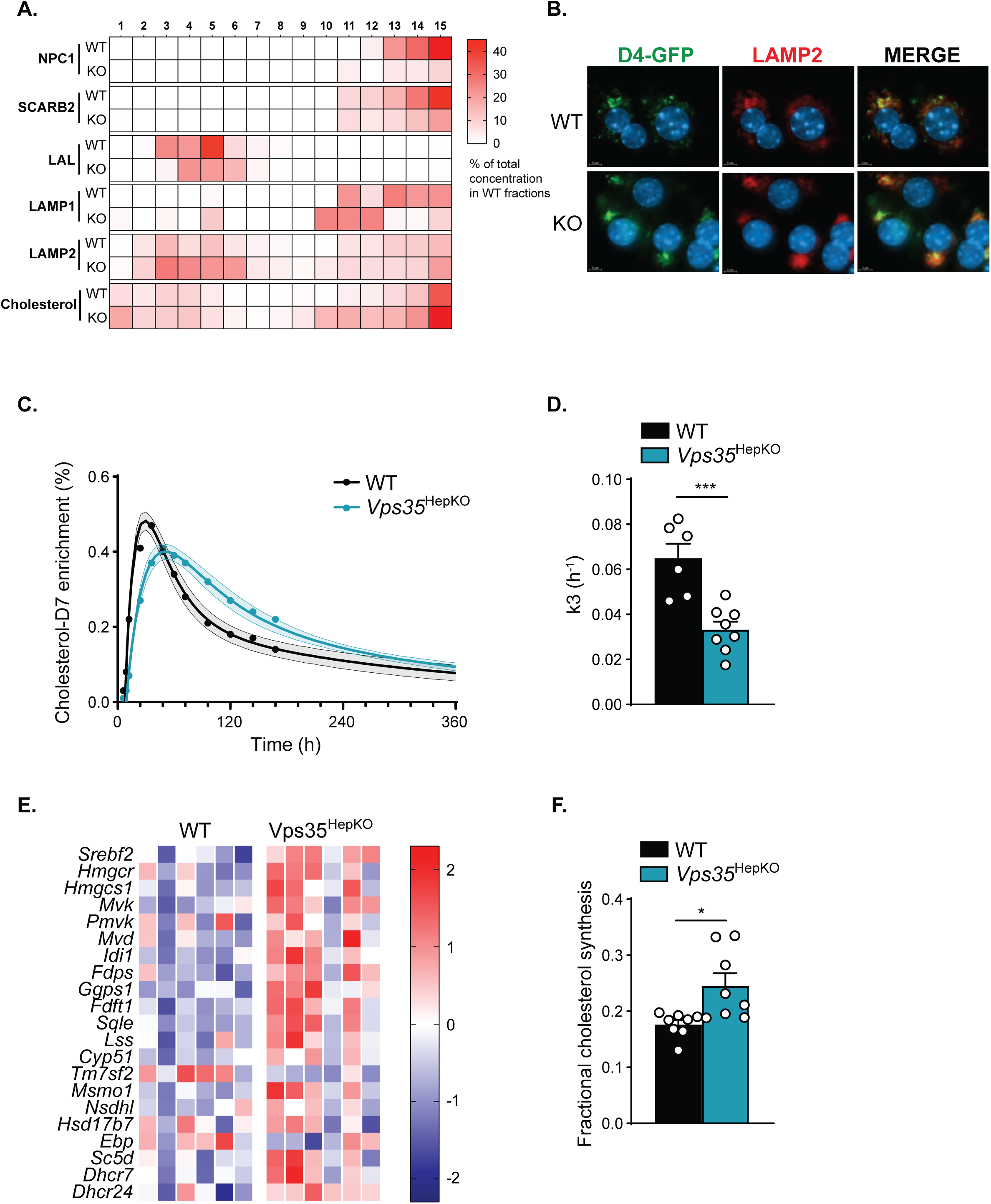
Hepatic secretion of endocytosed cholesterol is impaired, while cholesterol biosynthesis is increased in *Vps35*^HepKO^ mice. (**A**) Relative distribution of various lysosomal proteins involved in cholesterol handling within the endo-lysosomal system as determined by sucrose gradient fractionation and proteomics. Cholesterol levels in the sucrose gradient fractions of WT and *Vps35*^HepKO^ liver homogenate, determined by Amplex™ Red cholesterol assay. Protein and cholesterol levels are presented as the percentage of total levels in all WT fractions. (**B**) Cholesterol accumulation in WT and VPS35-deficient primary hepatocytes labeled with cholesterol probe GFP-D4 (green) and anti-LAMP2 antibodies (red). Nuclei were stained with DAPI. Scale bar 5 μm. (**C**) Fractional enrichment of orally administered cholesterol-D_7_ in WT and *Vps35*^HepKO^ mice (n=6-8). (**D**) Kinetic parameter *k3* describing the rate of cholesterol-D_7_ secretion by the liver into the circulation in WT and *Vps35*^HepKO^ male mice (n=6-8). (**E**) Heatmap presenting the z-score normalized expression of genes in the cholesterol biosynthesis pathway in livers of WT and *Vps35*^HepKO^ male mice (n=6). (**F**) Fractional cholesterol synthesis in WT and *Vps35*^HepKO^ mice (n=8). Data are presented as mean ± SEM; * p < 0.05 and *** p < 0.001.

To study the relative distribution of cholesterol, we measured the cholesterol content in the same fractions that were used for proteomics. We confirmed that VPS35 loss increased hepatic cholesterol levels (Figure 5A and S8A) and found that this increase was mainly seen in fractions positive for LAMP1 and LAMP2 (Figure 5A). To confirm that VPS35 deficiency leads to the accumulation of cholesterol in late endosomes/lysosomes, we specifically stained intracellular cholesterol with GFP-tagged version of the D4 domain of perfringolysin O (GFP-D4). This fluorescent probe can stain sterols in the plasma membrane and subcellular compartments such as lysosomes^45^. We observed that ablation of VPS35 in primary hepatocytes results in a strong perinuclear cluster of GFP-D4-positive compartments, which are partially co-stained with LAMP2, while in WT cells GFP-D4/LAMP2-positive compartments are distributed around the nucleus (Figure 5B). These data suggest that hepatic VPS35 loss results in cholesterol accumulation in late endosomes/lysosomes.

### Hepatic VPS35 deficiency increases hepatic cholesterol synthesis

We next investigated how the loss of lysosomal cholesterol metabolism-related proteins in *Vps35*^HepKO^ mice affects whole-body cholesterol homeostasis by studying cholesterol fluxes in *Vps35*^HepKO^ and control mice^29^. We administered D_7_-labeled cholesterol orally and measured the enrichment of this tracer in the blood for seven consecutive days (Figure 5C). The kinetic parameters of cholesterol-D_7_ turnover were calculated by curve fitting, and we found that the rate of cholesterol-D_7_ secretion by the liver into the circulation was markedly reduced in *Vps35*^HepKO^ mice compared to controls (Figure 5C,D). Yet, fractional intestinal cholesterol absorption was similar between groups (Figure S8B). These data indicate that after receptor-mediated endocytosis of D_7_-cholesterol-containing chylomicron remnants, hepatic secretion of chylomicron-derived cholesterol into the circulation is delayed in *Vps35*^HepKO^ mice compared to controls. This observation suggests that the transport of endocytosed cholesterol out of the lysosome to the ER is compromised upon hepatic *Vps35* ablation, which is in accordance with the observed lysosomal cholesterol accumulation (Figure 5A, B).

The observed differences in cellular cholesterol distribution may affect *de novo* cholesterol production. In NPC and LAL-D patient fibroblasts, and *Npc1*^-/-^ and *Lal*^-/-^ mice, it has been shown that impaired lysosomal cholesterol egress is associated with increased SREBP-mediated cholesterol biosynthesis in both NPC and LAL-D patient fibroblasts, and *Npc1*^-/-^ and *Lal*^-/-^ mice^4,56–58^. Consistently, we found using transcriptome analysis that the SREBP2-driven cholesterol biosynthesis is upregulated in *Vps35*^HepKO^ mice, with pronounced upregulation of genes such 3-hydroxy-3-methylglutaryl-CoA synthase 1 *(Hmgcs1),* mevalonate kinase *(Mvk)*, and squalene epoxidase *(Sqle)*, (Figure 5E). To assess the possible functional consequences of these observations, we studied *de novo* cholesterol synthesis. To this end, we gave mice drinking water containing the labeled cholesterol precursor ^13^C-acetate for 5 days and measured the incorporation of this tracer into cholesterol from bloodspots. Mass isotopomer distribution analysis (MIDA) was then applied to calculate fractional cholesterol synthesis^30,31^. We found that loss of hepatic VPS35 significantly increased whole-body cholesterol synthesis rates (Figure 5F), which predominantly reflects hepatic cholesterol synthesis in mice^58^. These data show that retromer plays an essential role in systemic cholesterol homeostasis.

### Vps35^HepKO^ mice display CE-enriched VLDL particles

A recent study showed that CESD is associated with CE-enriched VLDL particles^59^. These CEs are generated by hepatic sterol O-acyltransferase 2 (SOAT2; also known as ACAT2)^59^, an ER-resident enzyme responsible for cholesterol esterification, yielding CE for incorporation in VLDL particles^60,61^. Given that *Vps35*^HepKO^ mice show increased hepatic *Soat2* mRNA levels (Figure 6A), we explored whether this was associated with changes in SOAT2 levels and VLDL secretion and composition. Consistent with increased *Soat2* mRNA levels (Figure S9A), the hepatic SOAT2 levels were elevated (Figure S9B). Next, we determined the VLDL-TG secretion rate in mice by measuring plasma TG levels at different time points after injection of the lipoprotein lipase (LPL) inhibitor P407. Although we did not detect differences in VLDL secretion rate (Figure S9C), we found that the lipid composition of VLDL particles (phospholipids (PL): cholesterol: TGs, corrected for total ApoB levels) was different between control and *Vps35*^HepKO^ mice (Figure S9D). The VLDL particles of *Vps35*^HepKO^ mice had a relatively higher cholesterol content than in control mice (Figure S9D). Thus, hepatic loss of retromer results in elevated cholesterol-enriched VLDL particles, similar to those observed in patients with CESD.

**Figure 6.**
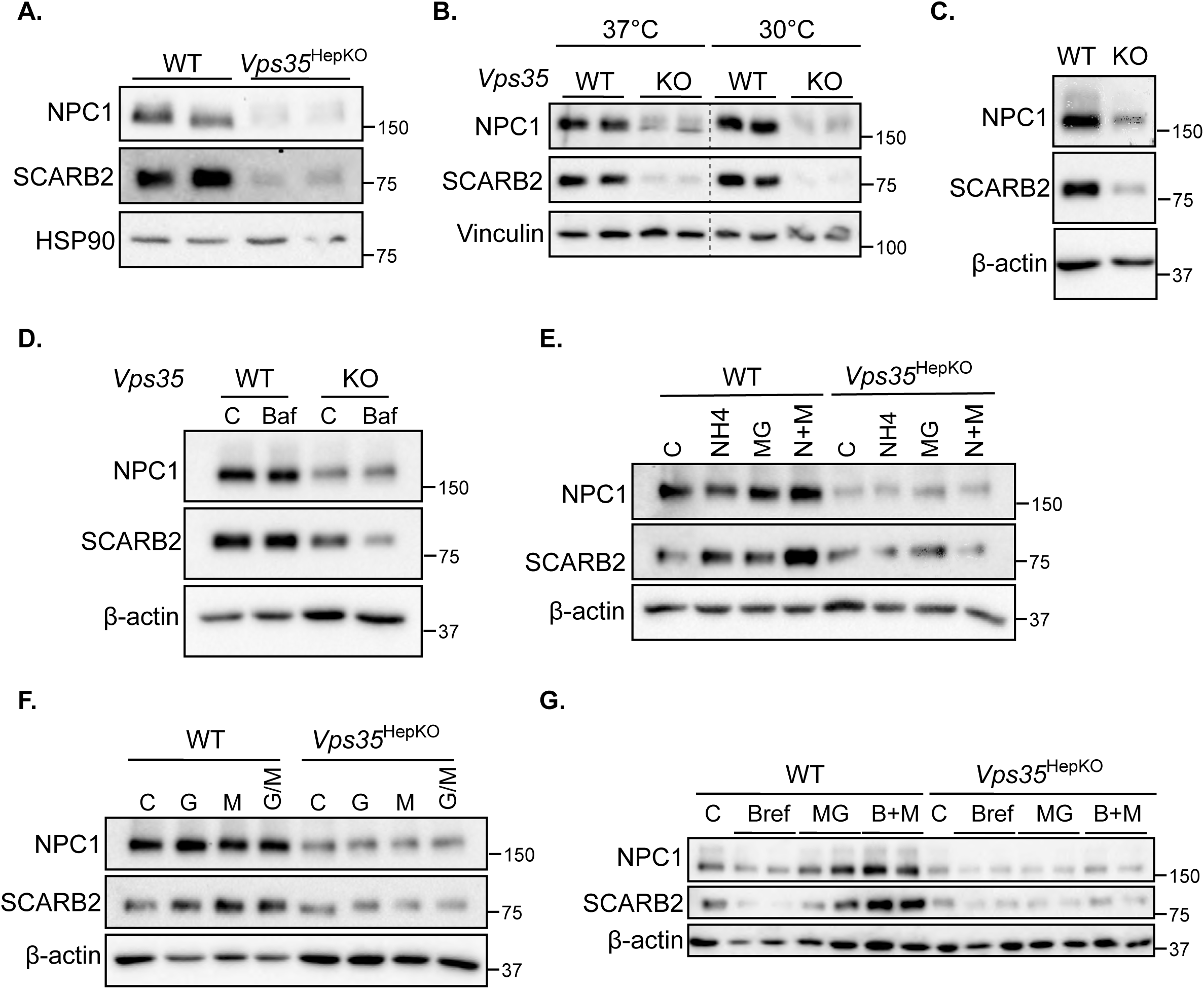
Reduced NPC1 and SCARB2 levels in *Vps35*^HepKO^ mice are not caused by protein misfolding, enhanced proteolysis, increased exosome release, or defects in the post-ER-Golgi trafficking pathway. (**A**) Protein levels of NPC1 and SCARB2 in primary hepatocytes isolated from WT and *Vps35*^HepKO^ mice (n=2). (**B**) Immunoblot of NPC1 and SCARB2 in WT and VPS35-deficient primary hepatocytes cultured at 37°C or 30°C. (**C**) Immunoblot of NPC1 and SCARB2 in precision-cut liver slices (PCLS) of WT and *Vps35*^HepKO^ mice, untreated or treated with (**D**) DMSO (C) or 100nM Bafilomycin A, (**E**) DMSO (C), 2.5 mM ammonium chloride (NH4), 5μM MG132 (MG), or 2.5 mM NH4 + 5μM MG for 16 h, (**F**) DMSO (C), 5 µM GW4869 (GW), 250 nM manumycin A1 (M), or 5 µM GW + 250 nM M (G/M) for 16 h, (**G**) 10 µg/mL Brefeldin A (Bref) (refreshing every 6 h), 2.5μM MG or 10 µg/mL Bref (refreshing every 6 h) + 2.5μM MG for 20 h.

### Reduced NPC1 and SCARB2 protein levels in Vps35^HepKO^ livers are not caused by post-translational mechanisms

To get mechanistic insight into how VPS35 loss leads to a decrease in NPC1 and SCARB2 levels, we studied cultures of primary hepatocytes isolated from *Vps35*^HepKO^ and control mice. We first confirmed that VPS35 deficiency blunts NPC1 and SCARB2 levels (Figure 6A), which again could not be explained by changes in mRNA levels (Figure S10A). A previous study showed that NPC1 can be degraded when it fails to be transported from the ER to late endosomes/lysosomes or is misfolded due to mutations^62^. NPC1 mutant proteins are either degraded via either ER-phagy, the selective ER degradation by autophagy, or ER-associated degradation (ERAD)^63^. To determine whether loss affects protein folding, we cultured primary hepatocytes of *Vps35*^HepKO^ and control mice at 30°C to improve the quality of protein folding^64^. However, lowering the temperature to 30°C did not improve NPC1 and SCARB2 protein expression compared to cells cultured at 37°C (Figure 6B).

To assess whether VPS35 deficiency promotes the proteolysis of NPC1 and SCARB2, we treated precision-cut liver slices (PCLS) from *Vps35*^HepKO^ and control mice with lysosomal inhibitors bafilomycin A1 or NH_4_Cl, and/or the proteasomal inhibitor MG132 for 16 h. We confirmed that VPS35 deficiency markedly reduces the protein levels of SCARB2 and NPC1 in PCLS that were cultured for 16 h as well (Figure 6C). However, the different inhibitors (as indicated in Figure 6D, E) that block the proteolysis of proteins could not rescue the strong reduction of NPC1 and SCARB2 levels. In contrast, the accumulation of ubiquitinated proteins was markedly increased in controls and *Vps35*^HepKO^ liver slices upon this treatment (Figure S10B, C), indicating that the used inhibitors were effective in blocking the proteolysis of proteins under these conditions. Next, we determined whether loss of VPS35 enhances the extracellular release of NPC1 and SCARB2 through exosomes. We treated the liver slices with GW4869 and/or manumycin A, which block the exosome biogenesis and release^65^. Similar to the results of the experiments with the proteolysis inhibitors, these two drugs did not reverse the reduction of NPC1 and SCARB2 levels in *Vps35*^HepKO^ liver slices (Figure 6F).

A recent study reported that loss of VPS35 in neuronal cells increases surface levels of NPC1 and SCARB2^55^. To assess whether this also occurs in livers, we determined the levels of NPC1 and SCARB1 at the PM, but in contrast to the previous study, we found a decrease in NPC1 and SCARB2 levels at the PM of VPS35-deficient liver cells compared to WT controls (Figure S10D).

To assess whether VPS35 deficiency affects NPC1 and SCARB2 post-ER trafficking and secretion, we treated control and *Vps35*^HepKO^ liver slices with Brefeldin A, an inhibitor that blocks the protein trafficking between the ER and Golgi apparatus, and thus protein secretion. Brefeldin A decreased NPC1 and SCARB1 levels in WT and *Vps35*^HepKO^ liver slices (Figure 6G), likely due to an augmentation of ERAD^63^. This further decrease in protein levels was normalized to the levels observed in untreated *Vps35*^HepKO^ liver slices by blocking proteasomal activity with MG132. In control liver slices, however, the levels of these two proteins were even increased after MG132 treatment (Figure 6D). These results indicate that the loss of VPS35 does not adversely affect post-ER trafficking or the cellular secretion of these proteins.

Taken together, these data suggest that the reduced levels of NPC1 and SCARB2 following VPS35 ablation are not caused by protein misfolding, enhanced proteolysis, increased exosome release, mislocalization to the cell surface, or defects in the post-ER-Golgi trafficking pathway.

## Discussion

Retromer is a central player in the endo-lysosomal network that facilitates the transport of cargos from endosomes back to the trans-Golgi network (TGN) or the PM, protecting them from lysosomal proteolysis^16–19^. Retromer dysfunction has been linked to neurodegenerative diseases such as Alzheimer’s (AD) and Parkinson’s disease (PD)^66–68^, and several studies have implicated retromer in cholesterol homeostasis^20,23^. However, its role in systemic cholesterol homeostasis remains unclear. To our knowledge, this is the first study showing that hepatic retromer is essential for maintaining cholesterol homeostasis. In particular, retromer is required to maintain the levels of several lysosomal proteins, including proteins such as LAL, SCARB2, and NPC1, that are key for lysosomal cholesterol egress, thereby regulating both hepatic and plasma cholesterol levels. In line with the phenotypes of LAL-D and NPC disease (MIM #613497, #620151, #278000, #257220, #607625), dysfunction of these proteins in VPS35-deficient livers is accompanied by CE and FC accumulation, likely in late endosomes/lysosomal compartments due to compromised CE hydrolysis and lysosomal cholesterol egress, respectively. Furthermore, NPC disease and LAL-D are associated with increased cholesterol biosynthesis^4,56–58^, a phenotype that we also observed in hepatic VPS35-deficient mice and is likely due to impaired movement of cholesterol out of the lysosomes to the ER, which results in enhanced activation of the SREBP2-activated cholesterol biosynthetic pathway.

The mechanism by which VPS35 depletion causes a strong reduction in LAL, SCARB2, and NPC1 levels, and whether the underlying mechanism is the same for these proteins, remains unclear. LAL is a lysosomal luminal protein, and modification of LAL with mannose-6-phosphate in the TGN is required for its efficient targeting to lysosomes as for many soluble lysosomal enzymes^69^. Retromer mediates the retrograde transport of CI-M6PR from the endosome back to TGN^21,22^, and since CI-M6PR is one of the two receptors responsible for the transport of M6P-modified proteins to the lysosomes^70^, loss of VPS35 may indirectly impair the transport of LAL from the TGN to the lysosomes. This defect in LAL sorting could result in increased extracellular LAL secretion, a phenomenon previously observed for cathepsin D^46,71–73^. Alternatively, the reduced LAL, NPC1, and SCARB2 levels may result from increased proteolysis, lysosomal exocytosis, or the release of extracellular vesicles (EV)^74^. Enhanced lysosomal exocytosis and exosome release are commonly observed under conditions of lysosomal dysfunction, likely to protect cells and maintain cellular homeostasis^74–78^. However, treatment of VPS35-deficient liver slices with inhibitors targeting lysosomal or proteasomal degradation pathways, or exosome biogenesis, failed to restore NPC1 and SCARB2 levels. Interestingly, blocking ER-to-Golgi and Golgi-to-PM transport with brefeldin A led to a further decrease in NPC1 and SCARB2 levels in both WT and VPS35-deficient liver slices, likely due to enhanced degradation via the ERAD system. This assumption is supported by experiments in which the combination of brefeldin A and the proteasomal inhibitor MG132 restored protein levels to those observed in the untreated WT and VPS35-deficient liver slices. Notably, while dual inhibition led to even higher protein levels in WT livers compared to untreated controls, this increase was not observed in VPS35-deficient liver tissue. Together with the observation that *Npc1* and *Scarb2* mRNA levels were not negatively affected in hepatic VPS35-deficient mice, it is plausible that retromer deficiency disrupts NPC1 and SCARB2 post-transcriptionally and before they are transported from the ER to the TGN, suggesting that loss of retromer impairs the translation of NPC1 and SCARB2. Notably, a recent study reported a novel link between VPS35 expression and translation. Here, they showed that loss of VPS35 reduces mitochondrial translation in HEK293T and lymphoblast cells by regulating the expression of the cationic amino acid transporter SLC7A1 at the plasma membrane^79^. However, as SLC7A1 is not expressed in the liver, these findings suggest that VPS35 regulates the translation of these lysosomal proteins through an alternative, yet unidentified, mechanism in hepatocytes.

Using H4 neuroglioma cells, Daly and colleagues also found a critical role for retromer in endo-lysosomal system^55^. They showed that retromer dysfunction causes mislocalization of various lysosomal membrane proteins, including LAMP1, LAMP2, and NPC1. In contrast to our findings, these proteins were primarily mislocalized to the cell surface in VPS35-deficient cells, but their total protein levels remained unchanged. In our study, in contrast, we did not observe increased redistribution of NPC1 and SCARB2 to the PM in VPS35-deficient livers. Instead, levels of both proteins were decreased at the PM compared to controls. Daly and colleagues also reported that VPS35 loss enhances lysosomal exocytosis of lysosomal lumen proteins, including LAL, NPC2 and multiple cathepsins (e.g., CATA, CATB, CATD, CATL), which they explained by lysosomal dysfunction. Similarly, we observed reduced total levels of LAL and CATD in VPS35-deficient livers but detected elevated NPC2 and CATB levels upon hepatic VPS35 ablation. Previous studies in NPC fibroblasts have demonstrated that NPC1 deficiency induces compensatory upregulation of NPC2^80–82^, which may explain the increased NPC2 levels in our model. The reason for the inconsistencies between Daly’s and our study remains unknown, but it could be explained by the different model systems and tissues that were examined.

Our study also reveals that *Vps35*^HepKO^ mice exhibit elevated plasma LDL cholesterol levels. In previous work, we showed that WASH and CCC, two multimeric protein complexes associated with retromer (Figure S1A,B), coordinate the endosomal recycling of LDLR and LRP1^9–11^. Interestingly, the current study suggests that retromer is not essential for endosomal trafficking of LDLR, but it plays a role in facilitating the LRP1 transport from endosomes to the PM, where it contributes to the uptake of chylomicron remnants, VLDL, and LDL. Retromer is known to recruit both the WASH and CCC complexes onto endosomes^25,83,84^, however, we did not observe any adverse effect on the levels or subcellular distribution of WASH and CCC components in *Vps35*^HepKO^ livers. These data strongly suggest that CCC/WASH-mediated endosomal LDLR transport is retromer-independent. A role for CCC/WASH in sorting of specific cargos, independent of retromer, has also been demonstrated in previous studies^24,25^. Given the lack of evidence for a role of retromer in the endosomal LDLR trafficking, our results suggest that the elevated plasma LDL cholesterol in *Vps35*^HepKO^ mice may instead result from increased hepatic cholesterol synthesis, likely due to impaired lysosomal cholesterol egress. We also found increased hepatic SOAT2 expression in *Vps35*^HepKO^ mice, which was linked with cholesterol-enrichment of VLDL particles ^60,61^, similar to what has been observed in patients with LAL deficiency^59^.

A recent study has associated retromer with metabolic-associated steatohepatitis (MASH) development and PCSK9 secretion^52^. In particular, it was found that loss of hepatic retromer reduces Sortilin-mediated PCSK9 secretion, resulting in elevated hepatic LDLR levels. Although we also observed increased hepatic LDLR protein levels in *Vps35*^HepKO^ mice, we did not find a decrease in plasma PCSK9. In fact, plasma PCSK9 levels were increased upon hepatic VPS35 depletion. We speculate that the increase in LDLR levels is a compensatory response to reduced LRP1 function in VPS35-deficient livers. This phenomenon has been well documented in previous studies investigating hepatic LRP1 function^49,85,86^. Interestingly, Ma and colleagues also showed that pharmacological stabilization of VPS35 reduced hepatic total and free cholesterol in a diet-induced MASH model^52^. While the mechanism underlying this cholesterol reduction was not addressed, our findings suggest that improved retromer function may enhance hepatic lysosomal cholesterol metabolism, potentially explaining the observed effect.

It is important to note that impaired retromer function has been associated with various neurodegenerative diseases. VPS35 levels are reduced in specific brain regions of patients with AD^68^, where retromer dysfunction has been linked to the accumulation of MAPT/Tau and amyloid beta (Aβ)^47,87,88^. Additionally, the *VPS35^D620N^* mutation is causative for an autosomal dominant form of PD^66,67^. Although it remains unclear whether this mutation disrupts systemic cholesterol homeostasis (e.g. hypercholesterolemia), several studies have reported that cholesterol homeostasis is often affected in neurodegenerative diseases^89–91^. These findings raise the intriguing possibility that retromer dysfunction may contribute to altered cholesterol homeostasis in the brain of AD and PD patients, beyond its well-established role in endosomal sorting of receptors and autophagy in neurons.

Taken together, our findings identify hepatic retromer as a key regulator of systemic cholesterol homeostasis. In addition to its well-established role in endosomal sorting of cargos, we uncovered that retromer is essential for maintaining the levels and function of key proteins in lysosomal cholesterol homeostasis, including LAL, SCARB2, and NPC1. While the mechanism remains to be fully elucidated, our results strongly suggest that loss of VP35 impairs the translation of NPC1 and SCARB2, highlighting a previously unknown role of retromer in mice and humans.

## Acknowledgements

The authors would like to acknowledge Niels Mulder, Desi Tsoneva, Trang Le, and Martijn Koehorst for their technical assistance with experiments, Ben Giepmans and Kim Kats for their assistance with EM, Philip Boucher for providing the GFP-D4 probe, and the lab of Peter Olinga for their help with precision-liver cut slice experiments. In addition, we thank Katharina Kuentzel and Dagmar Kratky for their valuable input and suggestions.

This study was financially supported by grants from the European Union (MSCA-ITN-2020, 953489; Acronym EndoConnect) to B.v.d.S, J.A.K., and J.H..; The Netherlands Cardiovascular Research Initiative: ‘the Dutch Heart Foundation, Dutch Federation of University Medical Centers, The Netherlands Organization for Health Research and Development and the Royal Netherlands Academy of Sciences’ (CVON2017-2020; Acronym Genius2) to J.A.K. and B.v.d.S.; ZonMW OC grant (2471032) to J.A.K. and B.v.d.S.; NWO ENW grants (OCENW.M.22.034 and OCENW.M.23.189) to B.v.d.S.; ZonMW OC grant (816729) to J.C.W. European Atherosclerosis Society Competitive grant and United for Metabolic Diseases (UMD) catalyst (2024-CG-004) grant to C.V.; the Graduate School for Drug Exploration (GUIDE); and the De Cock-Hadders Foundation to D.V. and M.G.B.; J.A.K. is an established investigator of the Netherlands Heart Foundation (2015T068). Electron Microscopy was performed in the UMCG Microscopy and Imaging Center (UMIC), sponsored by the Netherlands Organization for Scientific Research (ZonMW 91111.006; Ben Giepmans); and the Netherlands Electron Microscopy Infrastructure (NEMI), NWO National Roadmap for Large-Scale Research Infrastructure of the Dutch Research Council (NWO 184.034.014; Ben Giepmans).

## Contributions

D.Y.V., M.G.B., C.V., and B.v.d.S. conceived the study. D.Y.V., M.G.B., C.V., A.H.H., W.A.R.O., J.J.T., A.C., performed the experiments with technical support from M.S., N.H., N.J.K., A.G., Z.B., and R.H. M.H.K. performed histological stainings. A.R. assisted with the iPSC studies, and V.W.B assisted with the analyses of the different omics data. F.K. and J.F.d.B. assisted in the design of the cholesterol flux experiment, and J.F.d.B. analyzed the data. B.v.d.S., F.R., and M.M. analyzed the EM data. M.M.F., L.S., and J.H. performed lipidomics analysis. J.C.W. performed and analyzed the proteomics experiments. D.Y.V., J.A.K., C.V. and B.v.d.S acquired financial support. J.A.K. and B.v.d.S. supervised the work. D.Y.V., M.G.B., C.V., and B.v.d.S analyzed and interpreted data, and wrote the initial manuscript with all authors editing and approving the final text.

