## Supplemental info and data for "Hepatic retromer is essential for systemic cholesterol homeostasis by regulating lysosomal cholesterol metabolism"

#### Supplemental Figure 1

A.

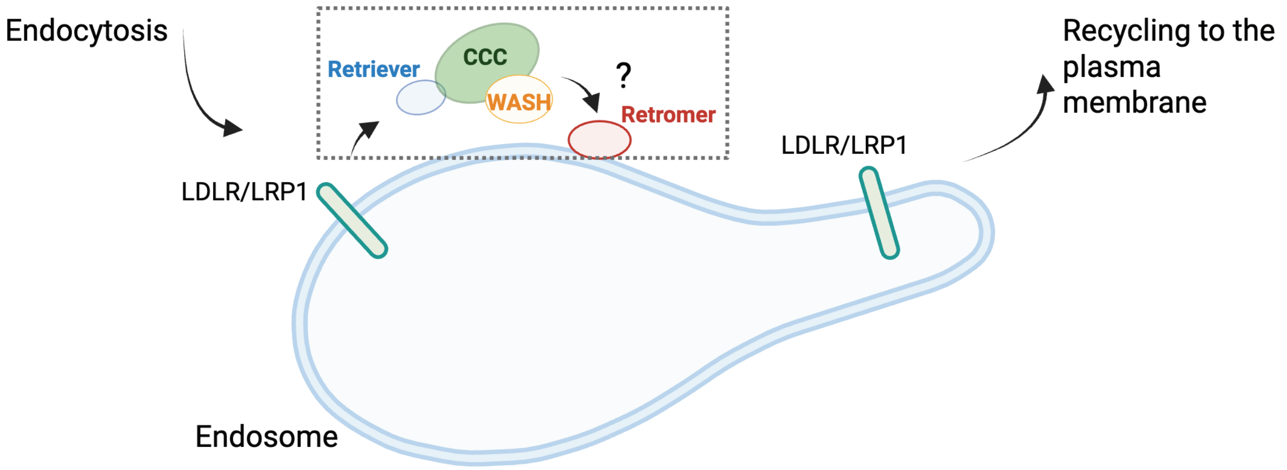


B.

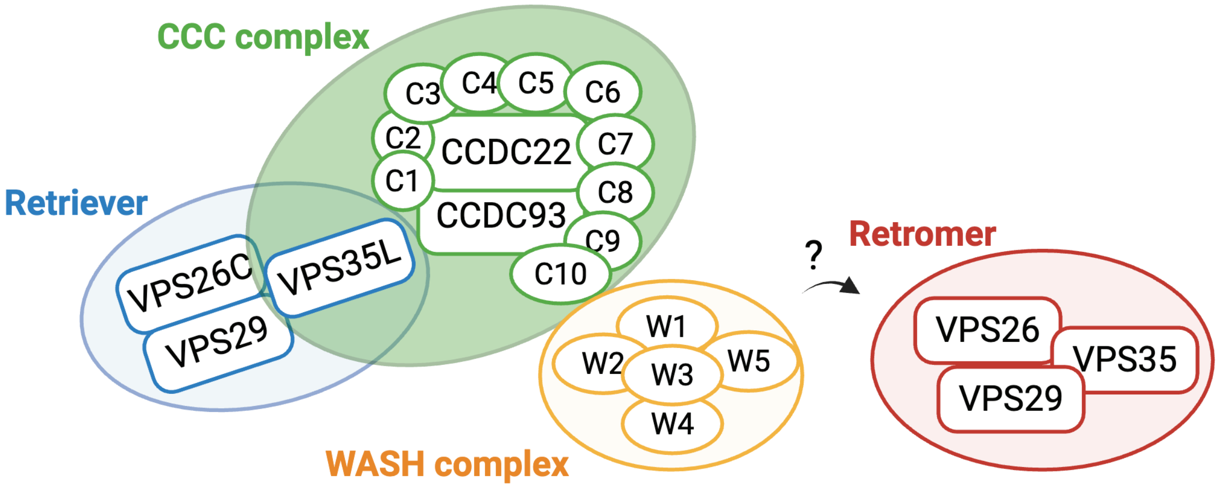

### Supplemental Figure 2

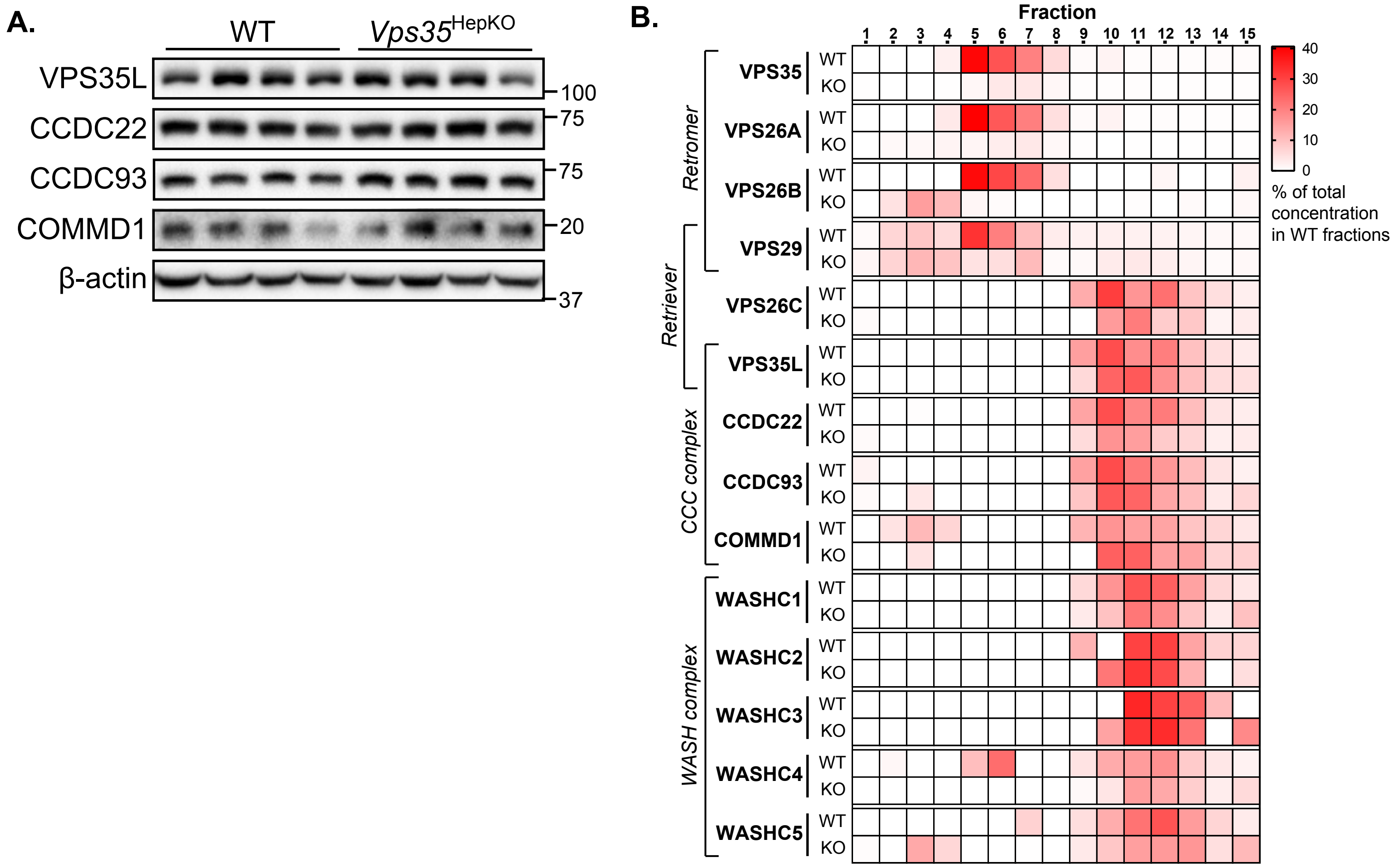

**Supplemental Figure 3**

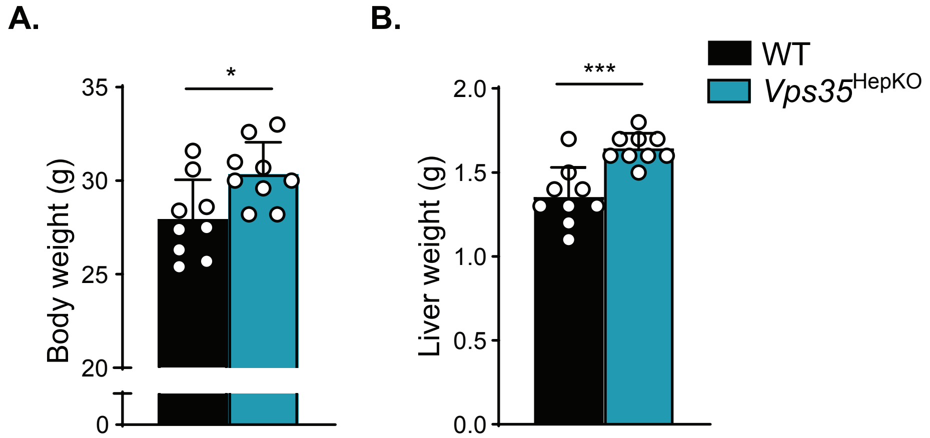

#### Supplemental Figure 4

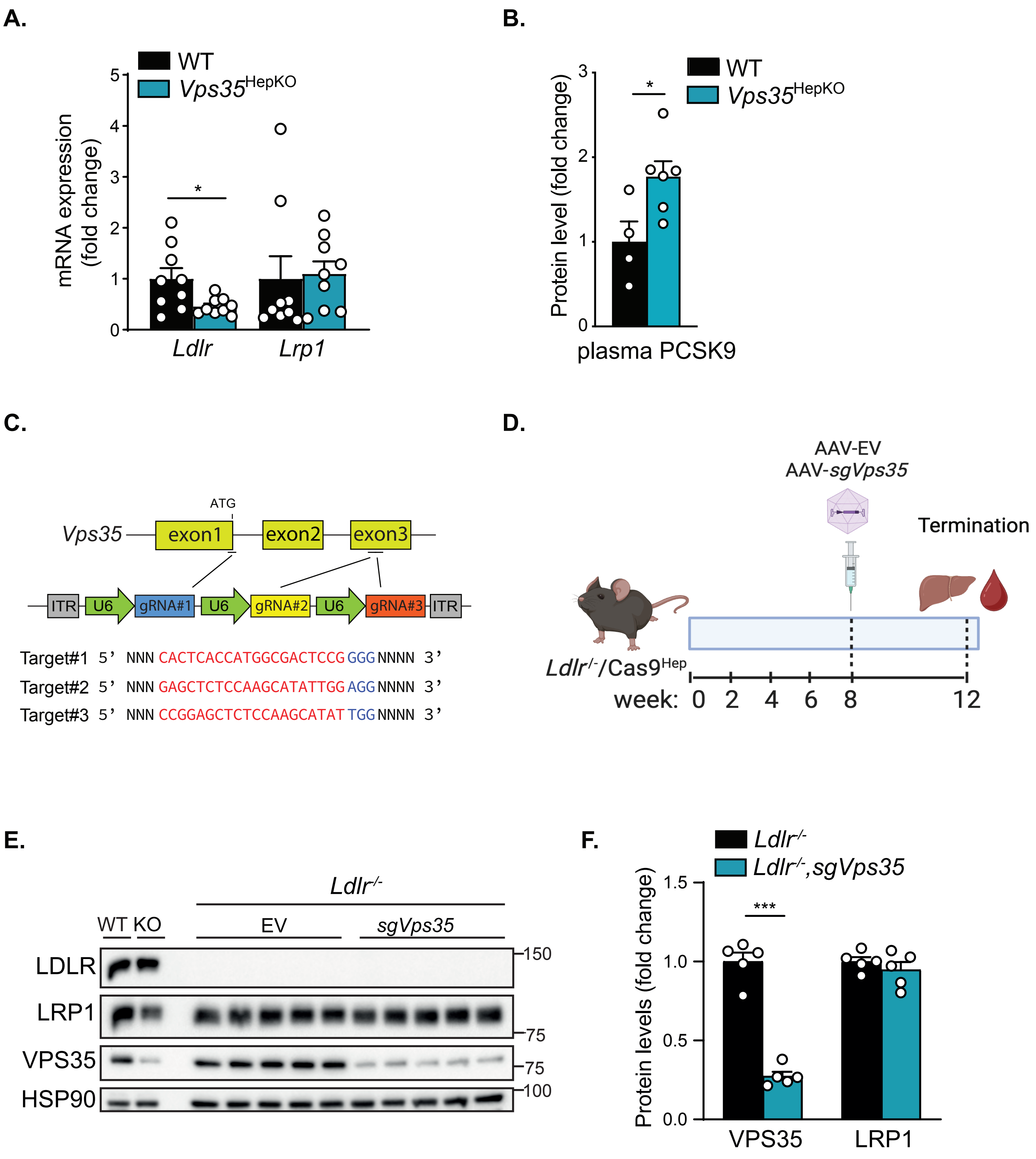

**Supplemental Figure 5**

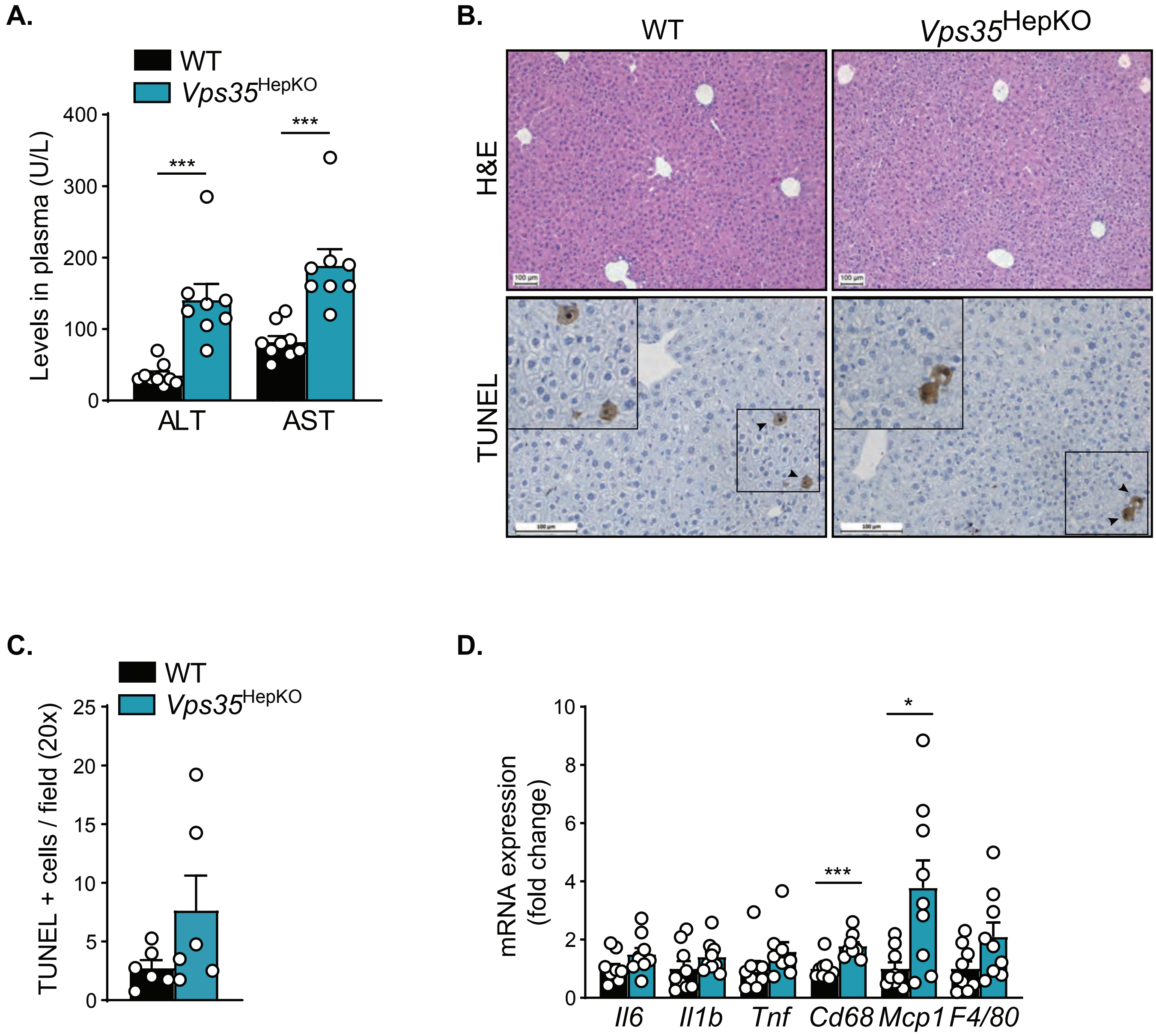

#### Supplemental Figure 6

A.

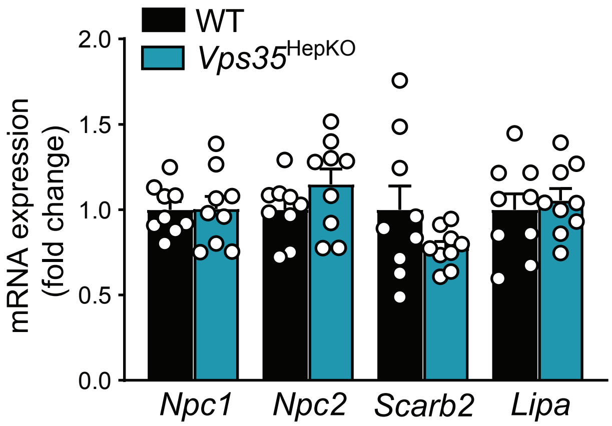


B.

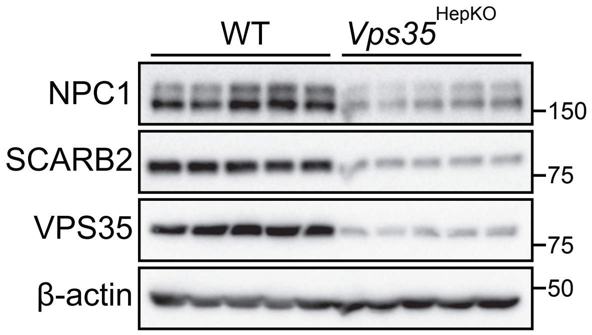


C.

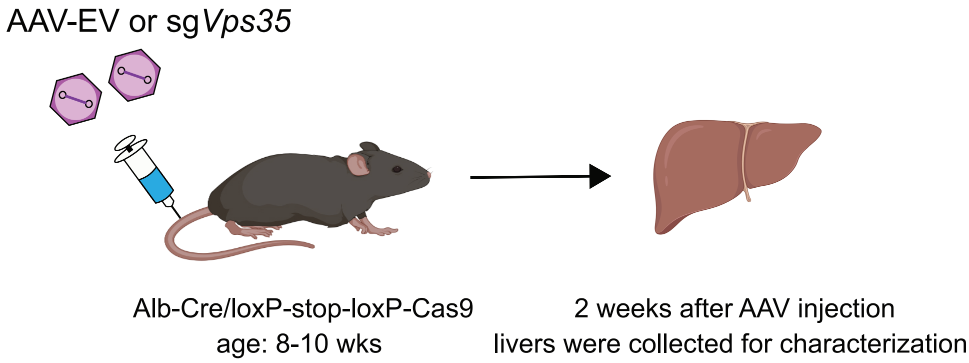


D.

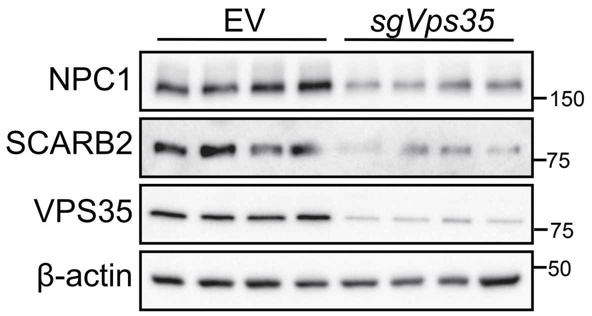

#### Supplemental Figure 7

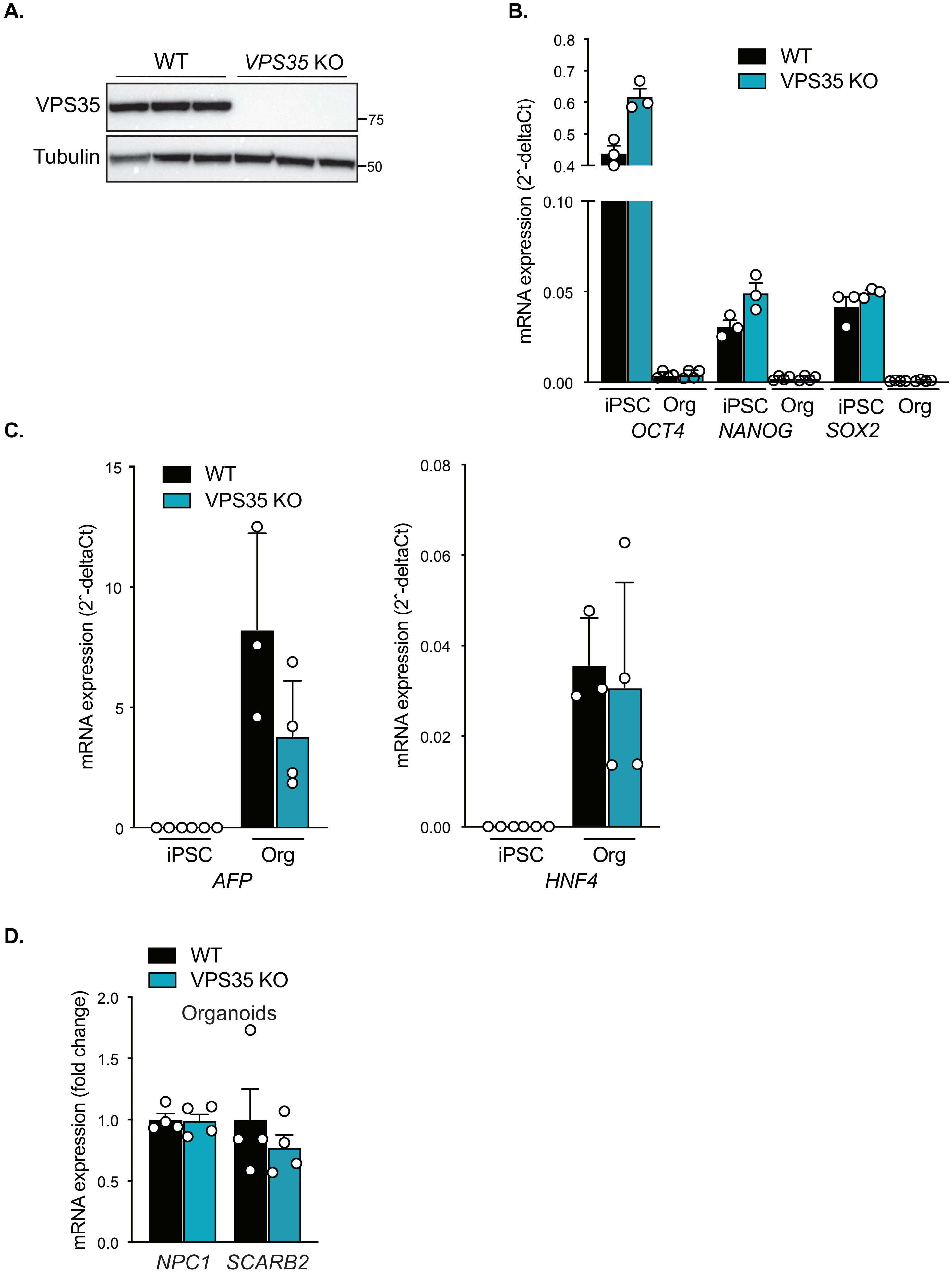

**Supplemental Figure 8**

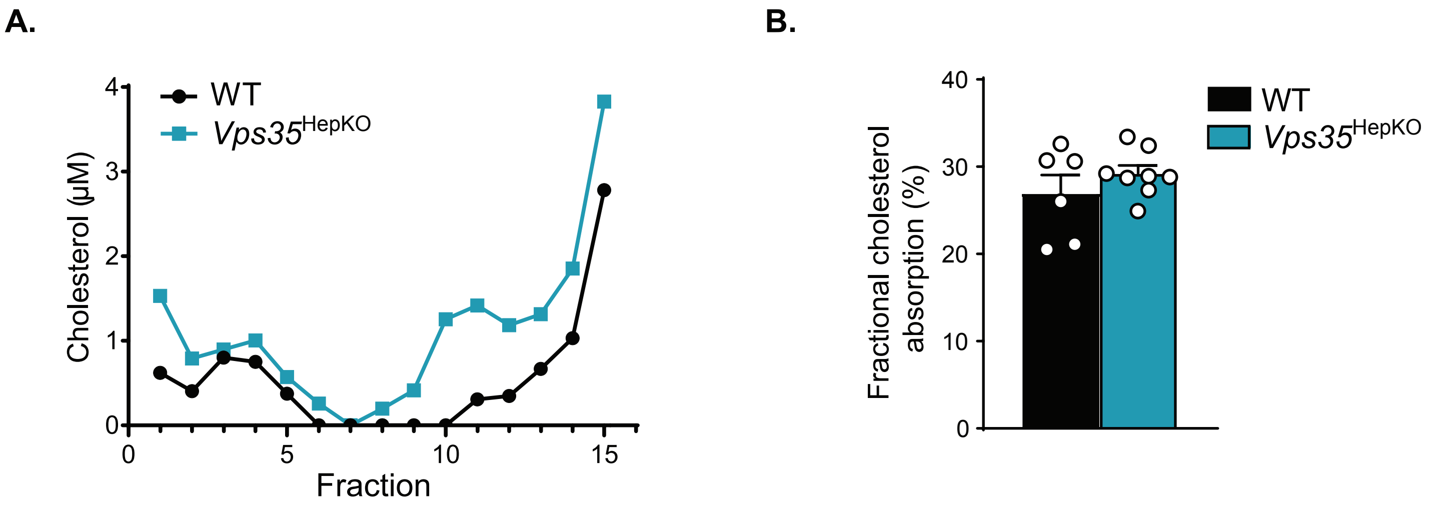

#### Supplemental Figure 9

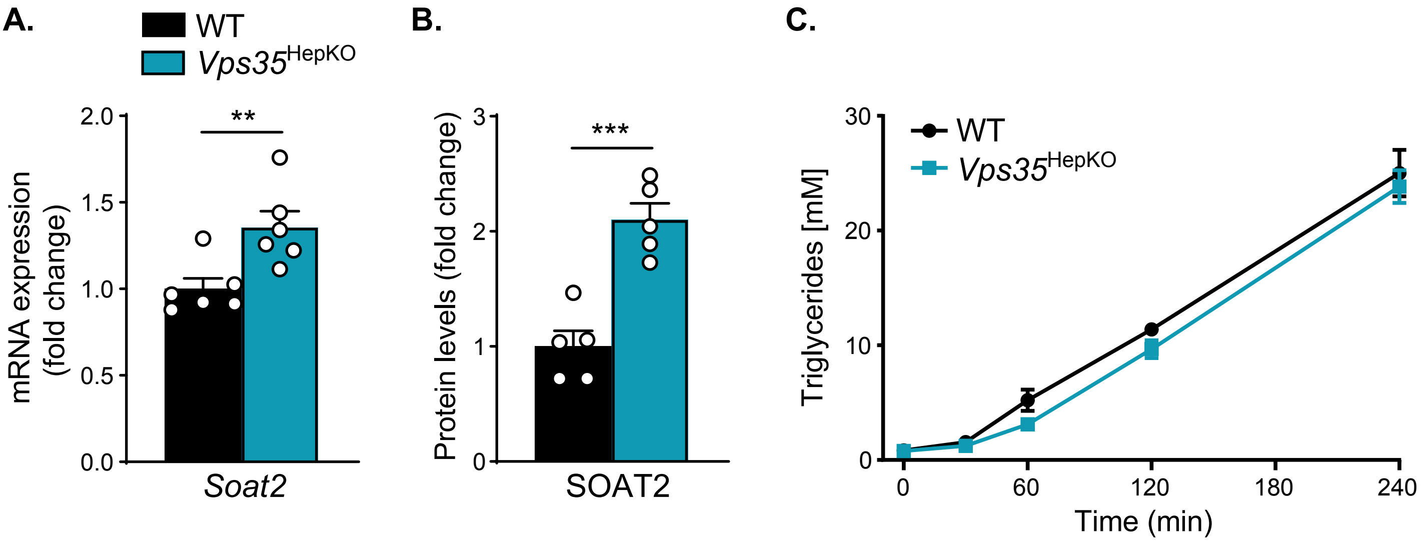


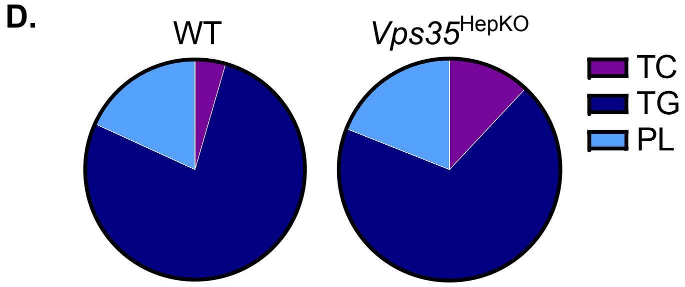


| | WT | *Vps35*HepKO |
|---|---|---|
| TC (%) | 4.5 ± 0.5 | 12.0 ± 1.4# |
| TG (%) | 77.2 ± 0.8 | 69.0 ± 2.8* |
| PL (%) | 18.3 ± 1.1 | 19.0 ± 1.8 |
| ApoB48 (%) | 85.2 ± 0.9 | 84.1 ± 0.5 |
| ApoB100 (%) | 14.8 ± 0.9 | 15.9 ± 0.5 |

#### Supplemental Figure 10

A.

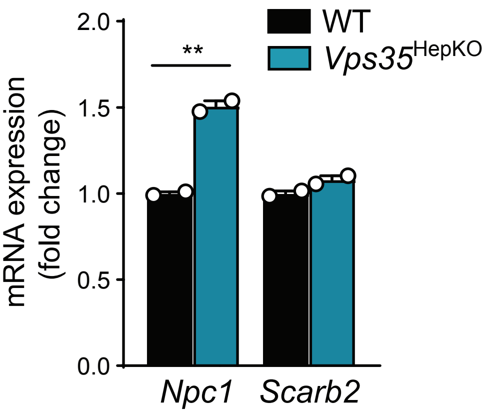


B.

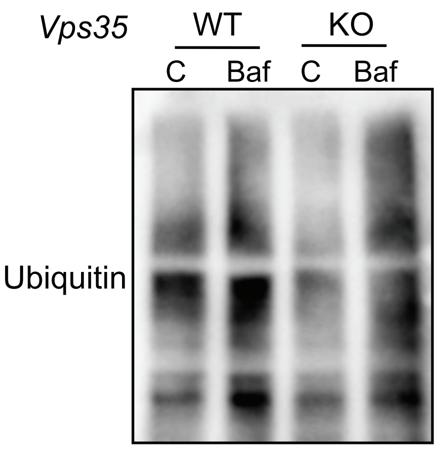


C.

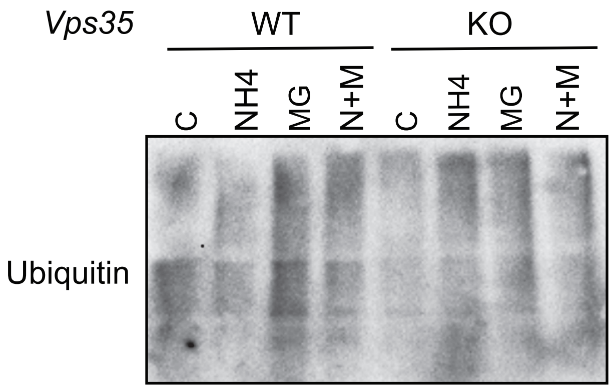


D.

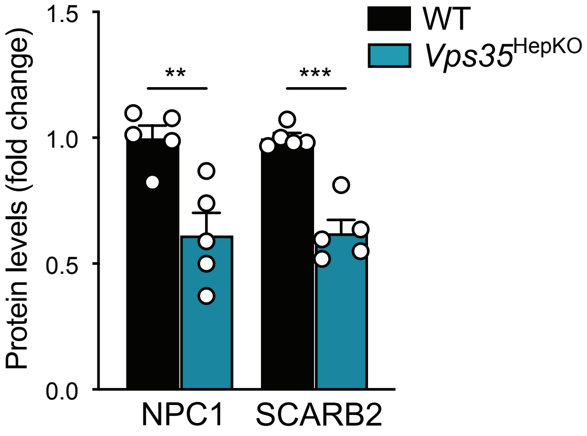

**Supplemental Figure Legends**

**Figure S1. Simplified model of the endosomal protein sorting complex. (A)** Upon endocytosis, receptors, like LDLR and LRP1, are recycled back to the cell surface. Our recent work demonstrates the essential role of the retriever, CCC, and WASH complexes in selective endosomal cargo sorting. However, the role of retromer in endosomal LDLR and LRP1 transport remains unclear. (**B**) The structure of the Commander complex with sub-assemblies, retriever and CCC complex, WASH complex, and retromer; subunits as C1-10 indicate COMMD proteins, W1-5 indicate WASHC proteins.

**Figure S2. Hepatic VPS35 deficiency does not affect the retriever, CCC and WASH complexes.** (**A**) Protein levels of retriever and CCC complex subunits in WT and *Vps35*$^{HepKO}$ male livers (n=4), determined by immunoblotting. (**B**) Relative distribution of retromer, retriever, CCC, and WASH subunits in WT and *Vps35*$^{HepKO}$ liver homogenates determined by sucrose gradient fractionation and proteomics; presented as the percentage of total protein levels of all WT fractions.

**Figure S3. Body and liver weight are increased upon hepatic *Vps35* ablation.** (**A**)**.** Body and (**B**) liver weights of WT and *Vps35*$^{HepKO}$ male mice at 18 weeks of age (n=9). Data are presented as mean ± SEM; * $p < 0.05$ and *** $p < 0.001$.

**Figure S4. Characterization of liver-specific VPS35-deficient mice and LDLR-deficient mice lacking hepatic VPS35.** (**A**) Relative mRNA levels of hepatic *Ldlr* and *Lrp1* in WT and *Vps35*$^{HepKO}$ male mice (n=9). (**B**) Plasma PCSK9 levels in WT and *Vps35*$^{HepKO}$ male mice, as determined by proteomics (n=5). (**C, D**) Schematic illustration of single-vector AAV (adeno-associated virus) system for targeting *Vps35* in (**D**) hepatic Cas9-expressing; LDLR-deficient mice (*Ldlr*$^{-/-}$;Cas9$^{Hep}$). Target sequences are shown in red and PAM sequences in blue. (**D**) Control and AAV-sgVps35 were injected in 8-week-old *Ldlr*$^{-/-}$;Cas9$^{Hep}$ mice. Four weeks after injection of the virus, mice were sacrificed. (**E**) Hepatic protein expression of LDLR, LRP1, and VPS35 in *Ldlr*$^{-/-}$;Cas9$^{Hep}$ mice and hepatic VPS35-deficient *Ldlr*$^{-/-}$;Cas9$^{Hep}$ male mice, determined by immunoblotting. The first two lanes are control liver homogenates from WT and hepatic VPS35-deficient mice, respectively. (**F**) Quantification of the protein levels in (**E**) (n=5). Data are presented as mean ± SEM; * $p < 0.05$ and *** $p < 0.001$.

**Figure S5. Hepatic loss of VPS35 does not lead to severe liver damage or liver inflammation.** (**A**) Plasma levels of ALT and AST (n=8-9). (**B**) Representative images of H&E and TUNEL staining of liver sections from WT and *Vps35*$^{HepKO}$ male mice; scale bars represent 100μm. (**C**) Quantification of TUNEL$^{+}$ cells from images shown in **B** (n=6). (**D**) Relative hepatic mRNA levels of inflammatory markers (n=9). Data are presented as mean ± SEM; * $p < 0.05$ and *** $p < 0.001$. *Abbreviations: ALT = alanine transaminase; AST = aspartate transaminase*

**Figure S6. Protein levels of NPC1 and SCARB1 in hepatic VPS35-deficient livers of female mice and in a model in which hepatic *Vps35* was acutely ablated** (**A**) Relative mRNA expression of *Npc1*, *Npc2*, *Scarb2,* and *Lipa* in WT and *Vps35*$^{HepKO}$ male livers (n=9). (**B**) Immunoblot of NPC1 and SCARB2 in WT and *Vps35*$^{HepKO}$ female livers (n=5), age 10 wks. (**C**) Somatic gene editing approach to study the acute effect of hepatic *Vps35* ablation on NPC1 and SCARB1 protein expression. (**D**) Protein levels of NPC1, SCARB2, and VPS35 in WT and *Vps35*$^{HepKO}$ male livers, 2 weeks after AAV injection (see **C**), Data are presented as mean ± SEM.

**Figure S7. Basic characterization of the human iPSCs and human iPSC-derived liver organoids.** (**A**) Protein expression of VPS35 in control WT and VPS35 knockout (KO) iPSCs, as determined by immunoblotting. (**B**) mRNA expression of several stemness and (**C**) liver organoids markers in WT and VPS35 KO iPSCs and liver organoids (WT and VPS35 knockout iPSC clones were differentiated into liver organoids, data presents 3-4 independent differentiation experiments). (**D**) mRNA expression of NPC1 and SCARB2 in WT and VPS35 KO human iPSC-derived liver organoids (n=4).

**Figure S8. Loss of hepatic VPS35 affects the distribution of hepatic cholesterol content, but not intestinal cholesterol absorption.** (**A**) Mean cholesterol concentrations determined by Amplex™ Red Cholesterol Assay in sucrose fractions of WT and *Vps35*$^{HepKO}$ liver homogenates. (**B**) Fractional intestinal cholesterol absorption in WT and *Vps35*$^{HepKO}$ mice (n=6-8), data are presented as mean ± SEM.

**Figure S9. Loss of hepatic VPS35 changes VLDL composition and lipoprotein receptor levels.** (**A**) Relative mRNA levels of hepatic *Soat2* in WT or *Vps35*$^{HepKO}$ male mice (n=6), determined by RNAseq analysis. (**B**) Relative SOAT2 protein expression in WT and *Vps35*$^{HepKO}$ livers (n=5), determined by proteomics. (**C**) Plasma TG levels (n=6-8) in WT or *Vps35*$^{HepKO}$ male mice, after i.p. injection with poloxamer 407. (**D**) VLDL particle composition in WT and *Vps35*$^{HepKO}$ male mice (n=3). TC, TG, and PL are presented as a percentage of total lipids corrected for total ApoB levels; ApoB48 and ApoB100 are presented as a percentage of total ApoB levels, * $p < 0.05$ and # $p < 0.01$. Except for (**D**), data are presented as mean ± SEM with ** $p < 0.01$ and *** $p < 0.001$.

**Figure S10. mRNA and protein levels of NPC1 and SCARB2 in WT and *Vps35*$^{HepKO}$ hepatocytes or plasma membrane-enriched fraction, respectively.** (**A**) Relative mRNA expression of *Npc1* and *Scarb2* in primary hepatocytes isolated from WT and *Vps35*$^{HepKO}$ mice (n=2). (**B**, **C**) The levels of ubiquitinated proteins in WT or *Vps35*$^{HepKO}$ liver slices after 16 hours of treatment with different inhibitors as indicated, determined by immunoblotting. C = DMSO, Baf = Bafilomycin (25 nM), NH4 = ammonium chloride (2.5 mM), MG = MG132 (2.5 μM), N+M = ammonium chloride (2.5 mM) and MG132 (2.5 μM). (**D**) Hepatic NPC1 and SCARB2 protein levels in plasma membrane-enriched fractions, determined by proteomics (n=5). Data are presented as mean ± SEM with ** $p < 0.01$ and *** $p < 0.001$.

#### Supplemental Tables

**Supplemental Table 1.** Sequence of the sgRNAs used in the current study.

| Gene | | Sequence (5'-3') |
|---|---|---|
| *Vps35* | sgRNA #1 | CACTCACCATGGCGACTCCG |
| | sgRNA #2 | GAGCTCTCCAAGCATATTGG |
| | sgRNA #3 | CCGGAGCTCTCCAAGCATAT |
| *VPS35* | sgRNA #1 | TACGAACTTGTACAGTATGC |
| | sgRNA #2 | GGTGTGCAACATCCCTTGAG |

**Supplemental Table 2.** Primer sequences used for qRT-PCR and PCR analysis.

| Gene | Forward primer (5'-3') | Reverse primer (5'-3') |
|---|---|---|
| *Cd68* | TGACCTGCTCTCTCTAAGGCTACA | TCACGGTTGCAAGAGAAACATG |
| *F4/80* | TGTGTCGTGCTGTTCAGAACC | AGGAATCCCGCAATGATGG |
| *Il1b* | TGCAGCTGGAGAGTGTGG | TGCTTGTGAGGTGCTGATG |
| *Il6* | CTGCAAGAGACTTCCATCCAGTT | AGGGAAGGCCGTGGTTGT |
| *Lipa* | TGTTCGTTTTCACCATTGGGA | CGCATGATTATCTCGGTCACA |
| *Ldlr* | TCCAATCAATTCAGCTGTGG | GAGCCATCTAGGCAATCTCG |
| *Lrp1* | GACCAGGTGTTGGACACAGATG | AGTCGTTGTCTCCGTCACACTTC |
| *Mcp1* | GCTGGAGAGCTACAAGAGGATCA | ACAGACCTCTCTCTTGAGCTTGGT |
| *Npc1* | TGTTTGGTATGGAGAGTGTGGA | GTCACAGCAGAGACTGACATTG |
| *Npc2* | AGGACTGCGGCTCTAAGGT | AGGCTCAGGAATAGGGAAGGG |
| *Ppia* | TTCCTCCTTTCACAGAATTATTCCA | CCGCCAGTGCCATTATGG |
| *Scarb2* | AGAAGGCGGTAGACCAGAC | GTAGGGGGATTTCTCCTTGGA |
| *Tnf* | GTAGCCCACGTCGTAGCAAAC | AGTTGGTTGTCTTTGAGATCCATG |
| *VPS35* | TAAATGGCTGACTGGGTGGA | GTGTTGTCGGTGCATCTCAA |
| *GAPDH* | AATCCCATCACCATCTTCCA | TGGACTCCACGACGTACTCA |
| *OCT4* | TGGGTGGAGGAAGCTGACAACAAT | TTCGGGCACTGCAGGAACAAATTC |
| *NANOG* | ATAGCAATGGTGTGACGCAGAAGG | CTGGTTGCTCCACATTGGAAGGTT |
| *SOX2* | CCTACTCGCAGCAGGGCACC | CTCGGCGCCGGGGAGATACA |
| *VPS35 exon1-2* | TGCCTACAACACAGCAGTCC | AAGTCCGGAGTTCACCAAGC |
| *VPS35 exon3-4* | CAGGCTGTGAAGGTCCAGTC | GTCCGGAGTTCACCAAGCATA |
| *ALB* | TTTATGCCCCGGAACTCCTTT | AGTCTCTGTTTGGCAGACGAA |
| *HNF4* | CACGGGCAAACACTACGG | TTGACCTTCGAGTGCTGCTCC |
| *AFP* | AGTGAGGACAAACTATTGGCCT | ACACCAGGGTTTACTGGAGTC |
| *SCARB1* | TGGGGATGCCTTCAAACAC | TTGAACTTCTGGGCAAATG |
| *LAL* | GAGAGCGGCCCGGCA | TGCAGGGTCCAGAGAACCAA |
| *NPC1* | AGCCAGTAATGTCACCGAAAC | CCGAGGTTGAAGATAGTGTCG |

**Supplemental Table 3.** Antibodies used for immunoblot analysis.

| Target | Company | Catalog # | Dilution |
|---|---|---|---|
| β-actin | Sigma | A5441 | 1:1000 |
| ApoB | EMD Millipore | 178467 | 1:1000 |
| Tubulin | Sigma | T9026 | 1:500 |
| CDCC22 | Proteintech | 16636-1-AP | 1:1000 |
| CDCC93 | Proteintech | 20861-1-AP | 1:1000 |
| COMMD1 | Proteintech | 11938-1-AP | 1:1000 |

| | | | |
|---|---|---|---|
| HSP90 | Cell Signaling | 4874 | 1:1000 |
| LAMP1 | BioLegend | 121602 | 1:1000 |
| LDLR | Abnova | PAB8804 | 1:1000 |
| LRP1 | Abcam | ab92544 | 1:1000 |
| NPC1 | Invitrogen | PA1-16817 | 1:1000 |
| SCARB2 | Cell Signaling | 27960 | 1:1000 |
| Vinculin | Cell Signaling | 4650 | 1:1000 |
| VPS26 | Abcam | ab23892 | 1:1000 |
| VPS29 | Abcam | ab98929 | 1:1000 |
| VPS35 | Abcam | ab10099 | 1:1000 |
| VPS35L | Thermofisher | PA5-28553 | 1:1000 |
| goat anti-mouse IgG (H+L)-HRP conjugate | Bio-Rad | 170-6516 | 1:10000 |
| goat anti-rabbit IgG (H+L)-HRP conjugate | Bio-Rad | 170-6515 | 1:10000 |
